# TDP-43-stratified single-cell proteomic profiling of postmortem human spinal motor neurons reveals protein dynamics in amyotrophic lateral sclerosis

**DOI:** 10.1101/2023.06.08.544233

**Authors:** Amanda J. Guise, Santosh A. Misal, Richard Carson, Hannah Boekweg, Daisha Van Der Watt, Thy Truong, Yiran Liang, Jen-Hwa Chu, Nora C. Welsh, Jake Gagnon, Samuel H. Payne, Edward D. Plowey, Ryan T. Kelly

**Affiliations:** Biogen Inc., Cambridge, MA 02142; Department of Chemistry and Biochemistry, Brigham Young University, Provo, UT 84602; Biology Department, Brigham Young University, Provo, UT 84602, USA

**Keywords:** TDP-43 proteinopathy, single cell proteomics, nanoPOTS, laser capture microdissection, amyotrophic lateral sclerosis (ALS), motor neuron, retromer complex, stathmin 2 (STMN2), human tissue pathology

## Abstract

Unbiased proteomics has been employed to interrogate central nervous system (CNS) tissues (brain, spinal cord) and fluid matrices (CSF, plasma) from amyotrophic lateral sclerosis (ALS) patients; yet, a limitation of conventional bulk tissue studies is that motor neuron (MN) proteome signals may be confounded by admixed non-MN proteins. Recent advances in trace sample proteomics have enabled quantitative protein abundance datasets from single human MNs (Cong et al., 2020b). In this study, we leveraged laser capture microdissection (LCM) and nanoPOTS (Zhu et al., 2018c) single-cell mass spectrometry (MS)-based proteomics to query changes in protein expression in single MNs from postmortem ALS and control donor spinal cord tissues, leading to the identification of 2515 proteins across MNs samples (>900 per single MN) and quantitative comparison of 1870 proteins between disease groups. Furthermore, we studied the impact of enriching/stratifying MN proteome samples based on the presence and extent of immunoreactive, cytoplasmic TDP-43 inclusions, allowing identification of 3368 proteins across MNs samples and profiling of 2238 proteins across TDP-43 strata. We found extensive overlap in differential protein abundance profiles between MNs with or without obvious TDP-43 cytoplasmic inclusions that together point to early and sustained dysregulation of oxidative phosphorylation, mRNA splicing and translation, and retromer-mediated vesicular transport in ALS. Our data are the first unbiased quantification of single MN protein abundance changes associated with TDP-43 proteinopathy and begin to demonstrate the utility of pathology-stratified trace sample proteomics for understanding single-cell protein abundance changes in human neurologic diseases.

## Introduction

Amyotrophic lateral sclerosis (ALS) is a progressive neurodegenerative disease that is commonly characterized by loss of spinal cord somatic motor neurons (MNs), muscle denervation, and loss of motor function. To date, approved pharmacologic treatments for ALS patients – riluzole, which targets glutamate toxicity; edaravone, which targets free radicals; and sodium phenylbutyrate/taurursodiol, aimed at reducing ER stress and mitochondrial dysfunction – have only very modest effects on disease progression (Miller et al., 2012; Paganoni et al., 2020; Witzel et al., 2022). Furthermore, biomarkers that indicate progression of the molecular cascade that leads to MN degeneration/loss remain a significant gap in the clinical development of investigational drugs. New molecular biologic insights are essential to meet the need of ALS patients.

Substantial human evidence points to a significant role for transactive response DNA-binding protein 43 (TDP-43) dysregulation in MN disease pathogenesis. *TARDBP*, the gene that encodes TDP-43, harbors C-terminal mutations in approximately 3% of patients with familial ALS (fALS) and sporadic ALS (sALS) (Sreedharan et al., 2008). Moreover, cytoplasmic inclusions of ubiquitinated, phosphorylated C-terminal TDP-43 fragments are a neuropathologic hallmark of sALS and most genetic subtypes of fALS (Arai et al., 2006; Hasegawa et al., 2008; Neumann et al., 2006). TDP-43 regulates transcription and mRNA splicing, stability, transport, and translation (Bjork et al., 2022). Transcriptomics studies have demonstrated that TDP-43 perturbations are associated with widespread alterations in mRNA expression and loss of cryptic exon suppression within specific transcribed regions (Fratta et al., 2018; Klim et al., 2019; Ling et al., 2015; Liu et al., 2019; Ma et al., 2022; Melamed et al., 2019). However, the contributions of individual cryptic exon inclusion events to development of ALS remain to be determined and the protein-level impacts associated with TDP-43 loss-of-function in MNs remain to be elucidated. Increased knowledge of protein-level dysregulation in motor neurons will augment our understanding of disease biology and has the potential to uncover new and cell type-specific biomarker candidates for ALS.

Proteomic studies of human ALS tissues have primarily focused on bulk tissue or pooled cell populations (Engelen-Lee et al., 2017; Hedl et al., 2019; Iridoy et al., 2018; Umoh et al., 2018); however, measurements of MN-relevant targets and cell-to-cell variability are can be obscured by cellular heterogeneity in bulk-tissue datasets. While single cell genomic and transcriptomic analyses benefit from amplification to boost signals from trace samples (Ho et al., 2020), protein-level characterization has remained challenging. Recent advances in trace sample proteomics have started to address these challenges, enabling the generation of quantitative protein abundance datasets from single motor neurons (Kelly, 2020).

Here, we present the first unbiased survey of protein expression dynamics in single human ALS spinal cord motor neurons sampled using laser capture microdissection (LCM) and analyzed with nanoPOTS (Zhu et al., 2018c) sample handling and ultrasensitive mass spectrometry-based detection. A pilot study of single MN proteome signatures readily revealed disease-state differentiability (ALS vs. control (CTL)) and demonstrated prominent abundance reductions in proteins that function in oxidative phosphorylation, mRNA splicing and translation, and retromer-mediated vesicular transport. In a follow-up experiment, we employed qualitative stratification of single MN proteome samples based on TDP-43 inclusion status. We found similar reductions in metabolic, RNA regulatory, and endolysosomal trafficking proteins in MNs with and without TDP-43 cytoplasmic inclusions. Our protein abundance datasets demonstrate reductions in *STMN2* as previously reported (Klim et al., 2019; Melamed et al., 2019), prominent reductions in endolysosomal trafficking complexes including the retromer and GARP complexes and elevated abundance of the neurexin CNTNAP2 in ALS MNs, all evident prior to the formation of detectable cytoplasmic TDP-43 inclusions. Against a backdrop of rapid advancements in single cell proteomics technologies, these experiments begin to demonstrate the potential for single cell proteomics to augment our understanding of motor neuron homeostasis and neuropathology in ALS.

## Results & Discussion

### Pilot Study - Single motor neuron proteomic profiling in human ALS spinal cord

We compared protein abundances in single thoraco-lumbar somatic motor neurons dissected from postmortem tissue sections from ALS donors (n=3) and control (CTL) donors (no history of neurologic disease; n=3). ALS tissue donors trended slightly older (range: 59-80 years) compared to the controls (range: 56-63 years). Postmortem intervals (PMIs) of ALS samples were slightly lower (range: 6.5-12 hours) compared to CTL samples (range: 13-15 hours). Individual ventral horn somatic MNs (n=6 per tissue donor) identified based on morphology and presence of Nissl substance were excised by LCM, captured into nanowells (Zhu et al., 2018b), and processed using the nanoPOTS workflow (Zhu et al., 2018c). Pooled samples of 10 MNs per donor were processed in parallel to single MN samples to improve Match Between Runs (MBR) (Zhu et al., 2018b). Samples were block randomized and analyzed by LC-MS/MS for protein identification and quantitation (**Figure 1A**). On average, ∼900 (µID=931±53; µQt=890±52; n=36) high confidence master proteins (HCMPs; 1% global FDR ∩ ≥ 1 unique peptide per protein) were detected in individual single ALS or CTL MNs (**Figure 1B**) and a total of 2515 proteins were identified across all samples (**Supplemental Table S1**). ALS and CTL MNs were readily differentiated by PCA-based dimensionality reduction of 1064 protein abundance measurements (**Supplemental Figure S1**). Acetylcholinesterase (ACHE) and peripherin (PRPH) were abundantly present in 67% and 100% of MN samples, respectively. Choline O-acetyltransferase (CHAT) was in low abundance and identified in only 22% of the single MN samples in this pilot (**Supplemental Table S1**). Moreover, MS-based CHAT detection was positive in 64% of CTL MN samples and 36% of ALS MN samples in our follow up experiment in which we selected MNs based on CHAT immunoreactivity in adjacent histologic sections (**Supplemental Table S3**). These findings are consistent with sub-optimal production or flyability of CHAT peptides and diminished ChAT expression in ALS MNs.

**Figure 1.**
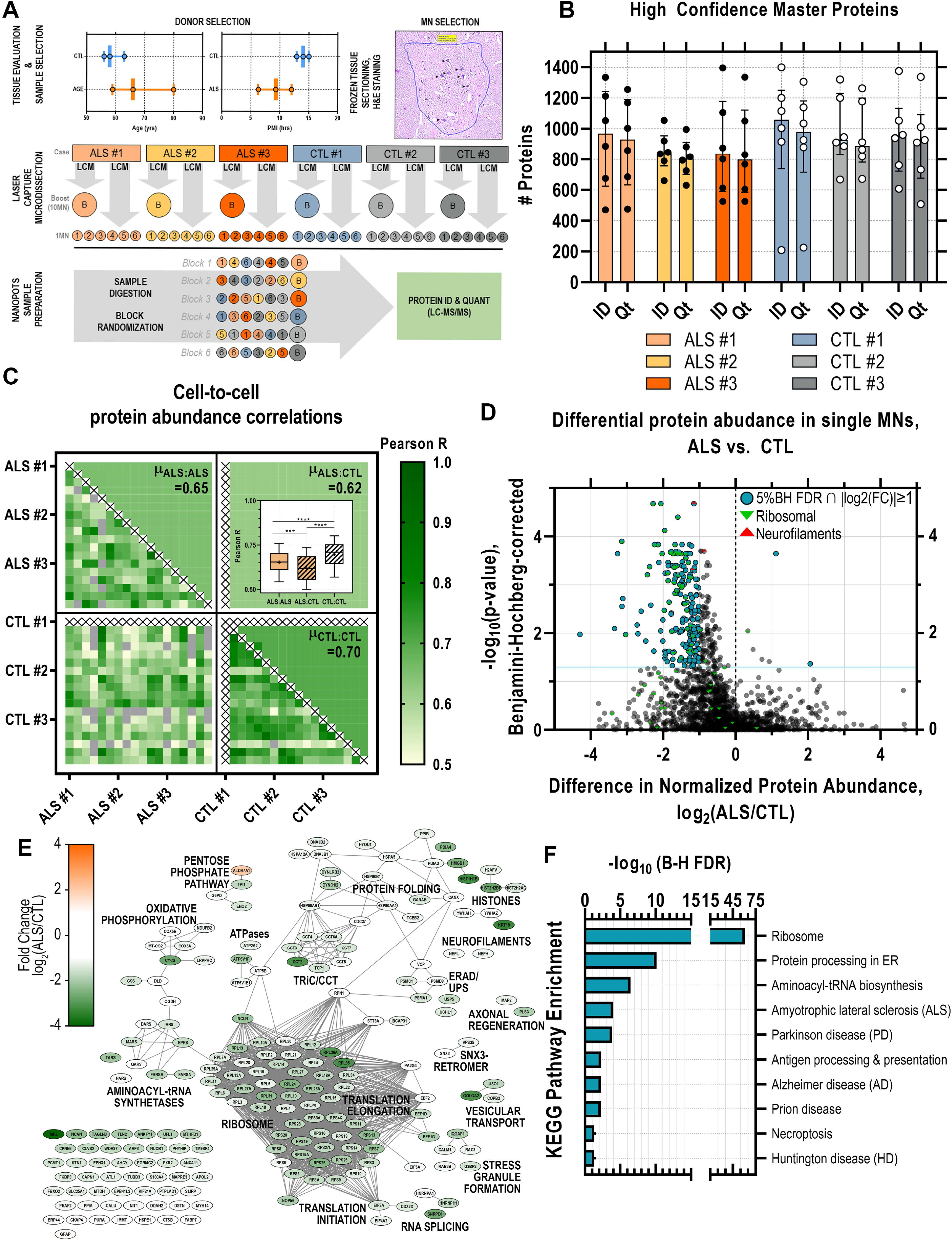
Ultrasensitive single cell proteomic mapping of ALS motor neurons. **(A)** Age and postmortem interval (PMI) of ALS and control (CTL) donor samples (n=3 per diagnosis) selected for single motor neuron (MN). Individual MNs (n=6 per donor) were identified in H&E-stained frozen tissue sections and excised by laser capture microdissection. Individual captured MNs were processed using the nanoPOTS workflow in parallel to “boost” samples (10 MN-equivalent) collected from each donor to facilitate match-between-runs. Samples were block randomized and analyzed by LC-MS/MS for protein identification and quantitation **(B)** High confidence (1% global FDR) Master Proteins (HCMPs) detected (ID) and quantified (Qt) across single MN samples isolated from ALS or CTL donor tissues **(C)** Pair-wise Pearson correlations between individual MN samples based on HCMP abundances and mean (µ) Pearson correlation score per comparison class (ALS:ALS, ALS:CTL, CTL:CTL) (*\*\*\*, p<0.001; ****, p<0.0001, 1-way ANOVA + Tukey’s MHC*) **(D)** Volcano plot of differential protein abundance (log_2_(ALS/CTL)) in ALS vs. CTL MNs based on normalized protein abundance values. Significantly dysregulated proteins (5% FDR (Benjamini-Hochberg-corrected p-value) ∩ |log_2_(ALS/CTL)|≥1) are indicated in blue; green marker overlays indicate ribosomal proteins; red marker overlays indicate neurofilaments. **(E)** Protein-protein interaction network of significantly differentially abundant proteins (5% FDR ∩ |log_2_(ALS/CTL)|≥1). Node color indicates fold change in normalized protein abundance (log_2_(ALS/CTL); edges denote high confidence STRING-db (v11.5) protein interactions (interaction score cutoff = 0.7) (F) KEGG pathway annotations significantly enriched among differentially abundant proteins. FDR reflects Benjamini-Hochberg-corrected p-value.

### Correlation scores between CTL:CTL single MN pairs were significantly greater than those of ALS:ALS single MN pairs

Within-group (ALS vs. ALS or CTL vs. CTL) and cross-group (ALS vs. CTL) comparisons to explore heterogeneity across single cells were performed by calculating Pearson correlation scores for all unique pairs of individual MNs (**Figure 1C**). The average pairwise correlation scores between ALS MNs (ALS:ALS) and between CTL MNs (CTL:CTL) were significantly higher (p<0.001 and p<0.0001, respectively; *1-way ANOVA with Tukey post-hoc test*) than pairwise correlation scores between ALS and CTL MNs (ALS:CTL) (**Figure 1C, *inset***), suggesting that even though individual MNs were collected from different ALS donors (n=3), variation between disease and healthy MNs appeared greater than the variation introduced from different donors. We also observed significantly higher correlation scores (p<0.0001, *1-way ANOVA with Tukey post-hoc test*) between pairs of CTL MNs relative to pairs of ALS MNs, which may suggest greater molecular heterogeneity among disease state MNs relative to their more homogenous healthy MN counterparts (**Figure 1C, *inset*).**

### Oxidative phosphorylation, mRNA splicing and translation, and retromer-mediated vesicular transport protein abundances are significantly reduced in ALS MNs

Quantitative comparison of 1870 normalized protein abundances from ALS and CTL MNs revealed 198 proteins significantly (*5% Benjamini-Hochberg FDR,* |log_2_(ALS/CTL)|≥1) altered in abundance, the vast majority (196/198) of which were decreased in ALS MNs relative to CTL MNs **(Figure 1D, Supplemental Table S2)**. In ALS MNs we observed reductions in neurofilament heavy (NEFH, FC=-1.18, *p=4.94E-04*), light (NEFL, FC=-1.15, *p=2.09E-05*), and medium (NEFM, FC=-0.87, *p=2.02E-04*) chain proteins relative to CTL MNs. Reduced neurofilament levels in ALS MNs could result from a variety of insults, including possible sequestration of neurofilament mRNAs within stress granules (Volkening et al., 2009). Neurofilament release, presumably following motor neuron death, can be measured as an ALS disease progression biomarker in CSF and plasma (Verde et al., 2021). The impact of the observed reductions in MN neurofilament levels on neurofilament biomarkers is unclear, however, studies that correlate neurofilament levels in tissues and fluids (Oeckl et al., 2020) could potentially refine our understanding of disease progression and therapeutic response biomarkers.

To further explore the relationships among proteins within the differentially abundant subset, we constructed a functional interaction network based on protein-protein interaction data curated by the STRING (v11) database (string-db.org) (**Figure 1E-F**). Diminished abundance of ribosomal proteins and translation factors is consistent with previous observations of translational dysregulation associated with ALS disease (Lehmkuhl and Zarnescu, 2018; Xue et al., 2020), while the resulting deficits in protein translation machinery could contribute to reduced global protein production in a feed-forward manner.

### Evaluating the broad dynamic range of protein detection in single human MNs and targeted investigation of proteins below the limit of detection

Comparison of median abundances measured for individual proteins identified in ALS and CTL MNs revealed sequencing of proteins spanning five orders of magnitude based on MS1 intensities, ranging from high-abundance structural neurofilament proteins (NEFH, NEFM, NEFL) to lower abundance membrane-associated synaptic proteins (LIN7C) **(Figure 2A)**. While downshifted in MS1 intensity, the relative protein rank orders in ALS MNs largely paralleled their counterparts in CTL samples, highlighting proteins at the lower end of the abundance range in CTL samples that fell below the limit of detection in ALS samples, such as Stathmin-2 (STMN2) **(Figure 2A)**. Presence-absence comparison of single cell proteomes identified 127 proteins uniquely present in CTL MN samples and 140 proteins uniquely present in ALS MN samples (**Figure 2B**). Interestingly, a substantial number of proteins detected in CTL MNs but absent in ALS MNs (**Figure 2B**, *teal*) are reported to be physical (**Figure 2C**, *network edges*) or indirect interacting partners (**Figure 2C**, *magenta node borders*) of TDP-43 (TARDBP, **Figure 2C**, *magenta node*), whose immunoreactive protein inclusions in neurons represent a distinguishing pathological feature of ALS (Mackenzie and Rademakers, 2008). An intriguing possibility is that sequestration of these interactors alongside TDP-43 within insoluble aggregates renders them less accessible for trypsin digestion – and subsequent detection – reflecting their functional unavailability. These interactors include multiple proteins with critical roles in mRNA processing and splicing (SNRNP200, NHP2L1, SRSF4, PRPF4, SART3, EIF4G3), stress-granule associated proteins (G3BP1, LSM12, LSM14A, FUS), and STMN2, a recently identified splice target of TDP-43 protein (Klim et al., 2019; Melamed et al., 2019). Following injury, STMN2 is recruited to axonal growth cones where it is proposed to function in repair and maintenance of the damaged axon, while loss of STMN2 has been proposed to accelerate degenerative programs in neurons and negatively impact NMJ stability (Graf et al., 2011; Shin et al., 2014; Shin et al., 2012). Aberrant splicing of STMN2 pre-mRNA has previously been reported in human spinal and cortical MNs from ALS donors, and suppression of STMN2 in iPSC-derived MNs resulted in inhibition of axonal regeneration in response to injury (Klim et al., 2019; Melamed et al., 2019). Orthogonal *in situ* hybridization-based detection of STMN2 RNA highlighted a significant decrease in abundance of STMN2 RNA in ALS MNs relative to CTL MNs **(Figure 2D-E)**, consistent with the diminished detection of STMN2 in single ALS MNs (**Figure 2A**). We further evaluated the presence of cryptic exon 2a-containing transcripts in adjacent tissue sections and demonstrate that single MNs expressing cryptic STMN2 exhibit low levels of canonical STMN2 (and vice versa) (**Figure 2F**), indicating that the loss of STMN2 protein observed by single cell proteomic analysis is likely the result of mis-splicing of STMN2. Interestingly, we noted clear loss of canonical STMN2 RNA expression also in MNs lacking apparent cytoplasmic TDP-43 inclusions (**Figure 2D**, *“ALS #7”, lower*), implicating functional impairment of TDP-43 in *STMN2* mis-splicing and deficiency prior to TDP-43-mislocalization and formation of histologically-evident inclusions.

**Figure 2.**
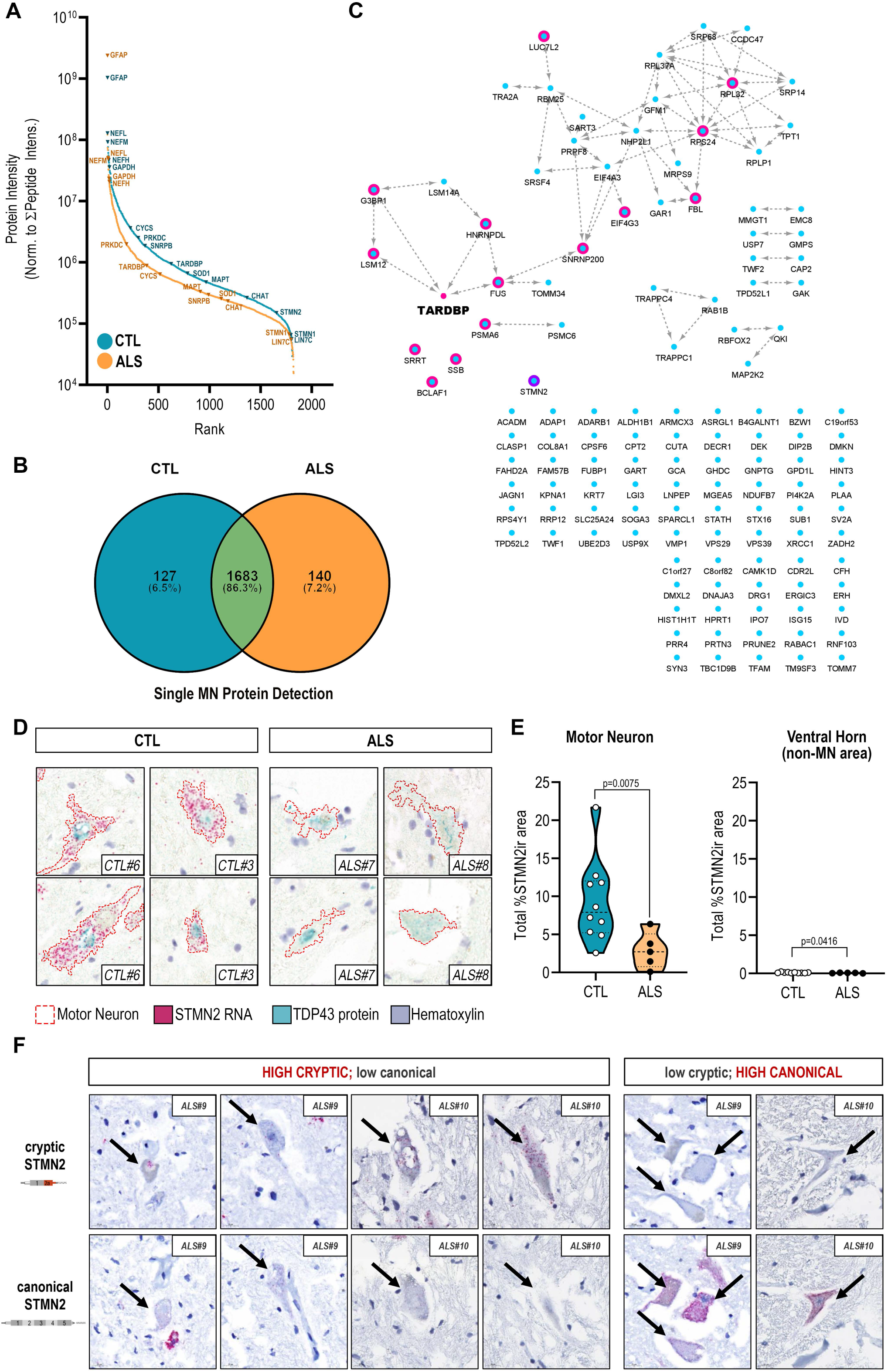
Absence of STMN2 protein detection in human ALS MNs parallels decreased abundance of Stathmin-2 (STMN2) RNA and absence of functional TDP-43 interaction partners. **(A)** Rank-ordered median MS1 intensities for proteins detected in either CTL (blue) or ALS (orange) single MNs span five orders of magnitude **(B)** Uniquely identified and overlapping proteins detected in ALS or CTL single MN samples **(C)** Functional interaction network of proteins uniquely detected in CTL single MNs and their relationship to TDP-43 (node manually added). Magenta node borders indicate reported interaction (direct or indirect) with TDP-43; purple node border indicates a downstream splice target of TDP-43, STMN2; edges indicate physical protein-protein interactions annotated by the STRING database (v11) **(D)** Dual expression of STMN2 RNA (red, ISH) and TDP-43 protein (teal, IHC) in human ventral motor neurons from ALS and CTL donors **(E)** Quantitation of STMN2 RNA signal in ventral MNs from ALS (n=5) and CTL (n=10) and donors as determined by relative percentage of STMN2-immunoreactive (STMN2ir) cell area; violin plots show total percent STMN2ir area in MNs and non-MN ventral horn tissue for each donor; *, p<0.05; **, p<0.01 unpaired t-test with Welch’s correction (F) Detection of cryptic exon-containing and canonical STMN2 transcript expression in adjacent ALS MN cross-sections.

### MN stratification by cytoplasmic TDP-43 inclusions for pseudotemporal assessment of protein abundance alterations

As postmortem CNS tissues often represent patients with end-stage ALS, we hypothesized that MNs at advanced stages of neurodegeneration could be overrepresented in our pilot experiment dataset. To test the hypothesis that MNs at an early stage of neurodegeneration might show a different proteomic profile compared to MNs at advanced stages of neurodegeneration, we generated a new expanded proteomics dataset with MNs stratified cells by the presence and extent of cytoplasmic TDP-43 inclusions **(Supplemental Figure S2)**. We employed TDP-43 immunohistochemistry in adjacent histologic sections **(Figure 3A)** to classify neurons in the following qualitative staging system: CTL = control MNs with normal appearing TDP-43; NON = ALS MNs with no overt cytoplasmic TDP-43 inclusions; MLD = ALS MNs with mild cytoplasmic TDP-43 inclusions; MOD = ALS MNs with moderate cytoplasmic TDP-43 inclusions; and SEV = ALS MNs with severe cytoplasmic TDP-43 inclusions (**Figure 3A, Supplemental Figure S2**). Single MNs were collected from three ALS and two CTL donors and target sample sizes per donor were determined based on *post-hoc* analyses of protein variability measurements across CTL MNs in the pilot dataset (**Supplemental Figure S3**) and models of sample size impacts on anticipated effect sizes for single cell proteomic data (Boekweg et al., 2021). We captured 25 MNs from each ALS donor, yielding between 13 and 28 MNs per stratified TDP-43 class (n_CTL_= 25; n_NON_= 15; n_MLD_= 28; n_MOD_= 13; n_SEV_= 18), resulting in a 33% increase in protein detection relative to our pilot study and comparison of 2238 proteins across TDP-43 strata. As in our initial TDP-43-agnostic analysis, we again observed evidence consistent with STMN2 depletion in ALS MNs. Decreased detection of STMN2 was apparent in ALS MNs lacking obvious cytoplasmic TDP-43 inclusions (NON) as well as MNs with TDP-43 cytplasmic inclusions (MLD, MOD, SEV) (**Figure 3B**), further suggesting that mis-splicing and diminished STMN2 translation are early events relative to the accumulation of cytoplasmic TDP-43 inclusions.

**Figure 3.**
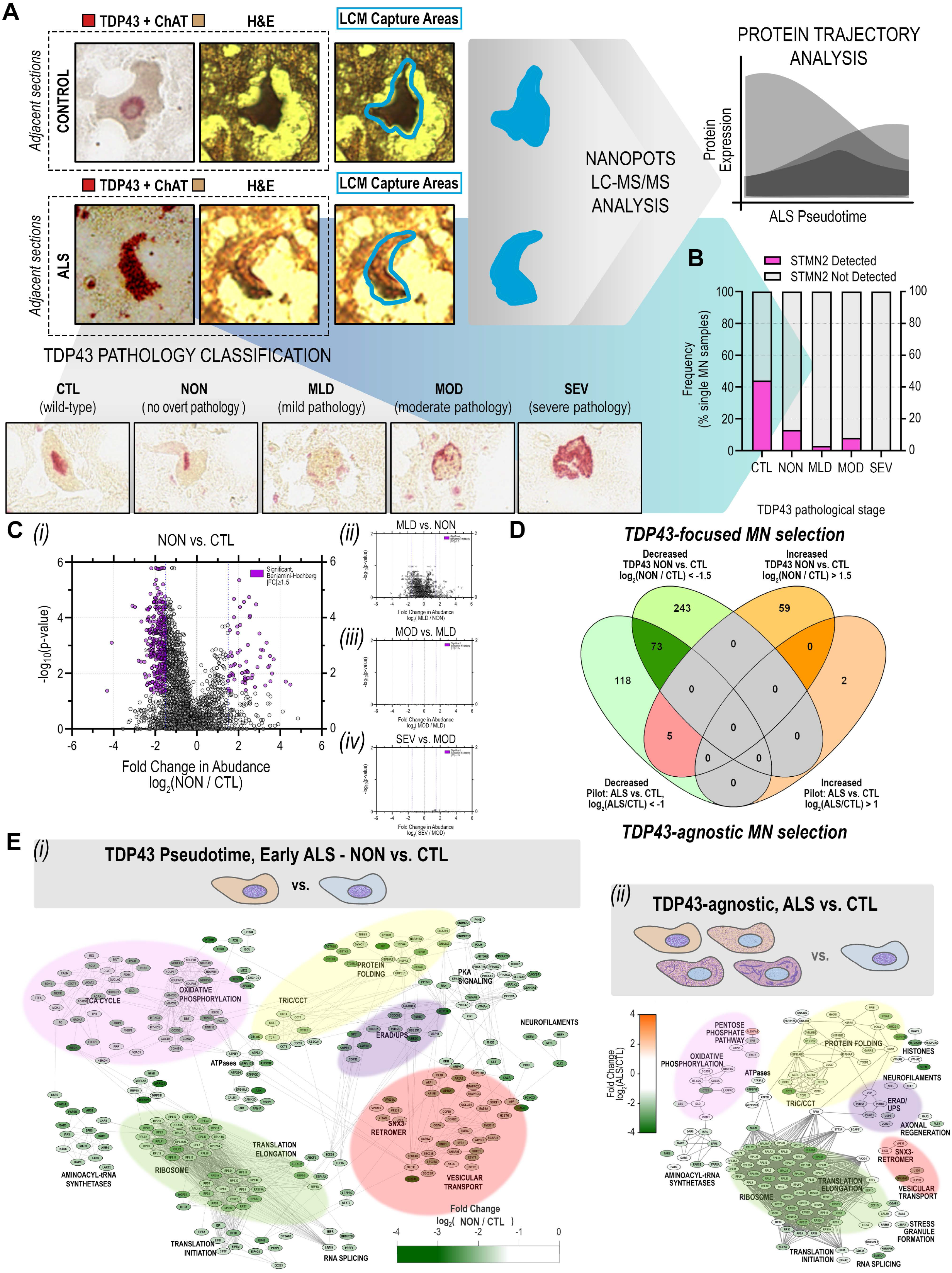
Significant disruption of proteostasis, mitochondrial dysfunction, and induction of pro-apoptotic signaling are apparent prior to overt TDP-43 aggregation. **(A)** Schematic of single MN selection for laser capture microdissection based on dual detection of TDP-43 and ChAT by immunohistochemistry in immediately adjacent human tissue sections. MNs selected for LC-MS/MS analysis by nanoPOTS were stratified based on the presence of TDP-43 inclusions (0, CTL – normal appearing; 1, ALS – normal appearing; 2, ALS – mild inclusions; 3, ALS moderate inclusions; 4, ALS – severe inclusions) for subsequent pseudotemporal comparisons **(B)** Frequency of STMN2 protein detection across single MNs corresponding to different TDP-43 inclusion strata **(C)** Volcano plot comparisons of differential protein abundance between sequential TDP-43 inclusion strata based on normalized protein abundance values (log_2_([Stage n+1]/[Stage n])). Significantly dysregulated proteins (5% FDR (Benjamini-Hochberg-corrected p-value) ∩ |log_2_([Stage n+1]/[Stage n]) ≥1.5) are indicated in purple *(i)* NON (ALS) vs. CTL (CTL); *(ii)* MLD (ALS) vs. NON (ALS); *(iii)* MOD (ALS) vs. MLD (ALS); *(iv)* SEV (ALS) vs. MOD (ALS) **(D)** Overlap in proteins designated as significantly differentially expressed in *(top)* “Early ALS” (NON vs. CTL, |log_2_([NON]/[CTL])| ≥1.5) following TDP-43-focused MN selection and in *(bottom)* ALS vs. CTL |log_2_([ALS]/[CTL])| ≥1.0) following TDP-43-agnostic cell selection **(E)** Overlap in protein complexes and functional pathways represented among the subset of proteins designated as significantly differentially expressed in *(i)* “Early ALS” (NON vs. CTL, |log_2_([NON]/[CTL])| ≥1.5) following TDP-43-focused MN selection or *(i)* in ALS vs. CTL |log_2_([ALS]/[CTL])| ≥1.0) following TDP-43-agnostic cell selection. Node color indicates fold change in abundance between groups; edges indicate high confidence protein-protein interactions curated by the STRING database (v11.5).

We observed dramatic and significant differential protein expression between healthy CTL MNs and ALS MNs lacking cytoplasmic TDP-43 inclusions (**Figure 3C*(i)***, CTL vs. NON) and, while not meeting the threshold for statistical significance set in this study, trends toward more subtle abundance alterations between normal-appearing ALS MNs and those exhibiting mild TDP-43 pathology (**Figure 3C*(ii),*** NON vs. MLD). Interestingly, we observed no significant stage-to-stage differences in protein abundances observed across MNs with greater complements of TDP-43 inclusions (i.e., MLD vs. MOD, MOD vs. SEV) (**Figure 3C*(iii, iv)*, Supplemental Figure S5**) **(Supplemental Tables S3-S6)**. We observed substantial overlap in differentially expressed proteins between ALS MNs in the TDP-43 agnostic pilot dataset (**Figure 1D-E, Supplemental Table S2**) and normal-appearing ALS MNs relative to CTL MNs (**Supplemental Table S5**), with 73 proteins significantly differentially expressed in both datasets, only 5 proteins showing opposite expression, and an overall trend toward protein deficiency in ALS MNs regardless of cytoplasmic TDP-43 inclusion status (**Figure 3D**). Differences in the subset of proteins identified in separate single cell experiments is unsurprising, due to the inherent stochasticity of data-dependent acquisition and the sparseness of single cell data in general, whose impacts and effects can further vary due to sample size. However, comparison of associated functions represented by individual proteins and protein groups across experiments showed clear deficiencies in proteins associated with oxidative phosphorylation, mRNA splicing and translation, and endolysosomal transport in normal-appearing ALS MNs with conservation of deficiency across TDP-43 inclusion strata (**Figure 3E**). Together, these data suggest marked changes in ALS MN proteomes *early* relative to the appearance of cytoplasmic TDP-43 inclusions.

### Endolysosomal sorting complex component abundances exhibit strong inverse correlations with increasing presence of cytoplasmic TDP-43 inclusions

Using the presence of cytoplasmic TDP-43 inclusions to stratify MN pathology, temporal profiles for individual proteins were determined based on Spearman correlation scores (R_s_) (**Supplemental Table S7**), resulting in the identification of a subset of proteins whose abundance values were either positively or inversely (**Figure 4A**) correlated (*q<0.05, Bonferroni*) with the extent of cytoplasmic TDP-43 inclusions per MN. We next looked among the subset of proteins exhibiting the strongest, negative correlations (≥95^th^ percentile; R_S_≥0.62) with increasing presence of cytoplasmic TDP-43 inclusions for over-representation of subcellular pathways (**Figure 4C*(i)***). In addition to neurodegenerative disease-associated KEGG pathway enrichments driven by the presence of metabolic (NDUFA7, NDUFV3, NDUFB8, MT-ND5, TIMM50) and proteasomal proteins (PSMC4, PSMD4, PSMD13, UCHL5), we observed significant over-representation of proteins associated with endocytosis, endolysosomal sorting, and retrograde endolysosomal trafficking, including: AMPH, AP2A2, CHMP4B, EHD3, SMAP1, SNX6, SNX12, VPS4B and VPS26B (**Figure 4C**).

**Figure 4.**
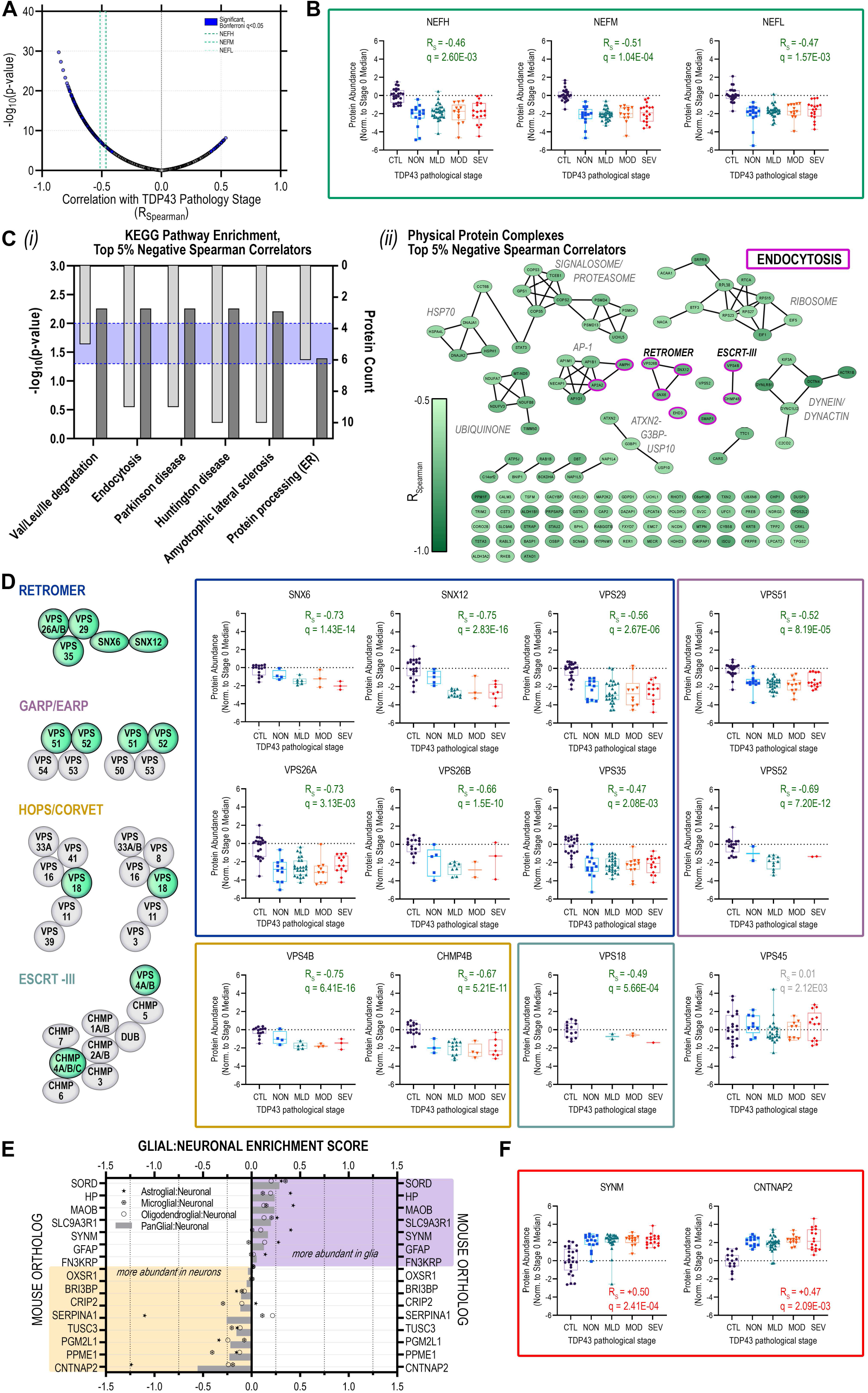
Neurofilament protein and retromer complex component abundances track inversely with increasing TDP-43 aggregation in postmortem human MNs. **(A)** Volcano plot of Spearman rank correlations between individual protein abundances per single MN and corresponding TDP-43 stratum; purple highlight, significantly (Bonferroni q < 0.05) correlated proteins; green dashed lines indicate neurofilament protein correlations (heavy (NEFH), medium (NEFM), light (NEFL)) **(B)** Distribution of protein abundances in individual MNs across TDP-43 strata for neurofilament proteins (NEFH, NEFM, NEFL) **(C)** (i) KEGG pathways over-represented and *(ii)* physical protein complexes represented among proteins comprising the top 5% of negative correlators with respect to TDP-43 strata; **(D)** TDP-43-inclusion-strata-associated protein abundance trajectories for retromer complex components SNX6, SNX12, VPS26A, VPS26B, VPS29, VPS35; GARP/EARP complex components VPS51 and VPS52; HOPS/CORVET complex proteins VPS18; and ESCRT-III complex proteins VPS4B and CHMP4B **(E)** Glial:Neuronal enrichment scores for proteins significantly positively correlated with increasing TDP-43 inclusion strata determined based on log_2_-transformed LFQ ratios between individual glial and neuronal populations measured in (Sharma et al., 2015). **(F)** TDP-43 pathology-associated protein abundance trajectories for the intermediate filament protein SYNM and the presynaptic neurexin protein CNTNAP2. Box plots indicate median ± IQR; whiskers indicate full data range per stage.

Separation of the strongest negative correlators with respect to increasing apparent severity of TDP-43 inclusions based on physical protein interactions revealed core components of the retromer (VPS26B/SNX6/SNX12), ESCRT-III (<u>e</u>ndosomal <u>s</u>orting <u>c</u>omplexes required for transport; VPS4B, CHMP4B), and GARP/EARP (<u>G</u>olgi- and <u>e</u>ndosome-<u>a</u>ssociated retrograde <u>p</u>rotein; VPS52) complexes (**Figure 4C*(ii)*)**. Remaining members of the heteropentameric retromer complex: VPS26A, VPS29, and VPS35; components of the CORVET/HOPS (<u>c</u>lass <u>C co</u>re <u>v</u>acuole/<u>e</u>ndosome tethering/<u>ho</u>motypic fusion and <u>p</u>rotein <u>s</u>orting) tethering and sorting complexes (VPS18) and GARP/EARP (<u>G</u>olgi-<u>a</u>ssociated retrograde <u>p</u>rotein/<u>E</u>R-<u>a</u>ssociated retrograde <u>p</u>rotein) transport complexes (VPS51, VPS52) also showed strong inverse correlations with increasing extent of cytoplasmic TDP-43 inclusions (**Figure 4D**).

Conserved across eukaryotic cells, the retromer complex recycles transmembrane protein cargoes from endosomes to the plasma membrane/cell surface, trans-Golgi network, or lysosomes (Vagnozzi and Pratico, 2019)). A key component of retromer cargo recognition, VPS35 dysregulation has been linked to the pathogenesis of Alzheimer disease (Filippone et al., 2021; Qureshi et al., 2022; Simoes et al., 2021; Wen et al., 2011), and Parkinson disease (Eleuteri and Albanese, 2019; Vilarino-Guell et al., 2011; Zimprich et al., 2011). Neuronal accumulation of insoluble phosphorylated TDP-43 was reported in conditional Vps35-knockout mice (Tang et al., 2020). Moreover, VPS35 mRNA-level deficiencies were documented in human cortical tissues from a cohort of donors harboring progranulin (PGRN) mutations and with FTLD (frontotemporal lobar degeneration) diagnoses (Tang et al., 2020). Several reports have linked retromer and/or endosome dysfunction to human ALS and/or disruption of TDP-43 proteostasis and cytoplasmic aggregates (Liu et al., 2017; Muzio et al., 2020; Shao et al., 2022). While the mechanism linking retromer/endosome deficiency with the accumulation of TDP-43 aggregates remains undetermined, one possible factor is impaired autophagic clearance of TDP-43 aggregates (Barmada et al, Scotter et al.) since autophagosome maturation is dependent on the endolysosomal pathway. Together, these data fit within a model in which TDP-43 protein levels are regulated both through proteosome-mediated and endolysosomal degradation in healthy MNs (**Figure 5A**). Upon disruption of protein sorting in the early/sorting endosome resulting from deficiencies in retromer complex proteins occurring in the early stages of ALS MN degeneration, TDP-43 cargoes are no longer efficiently trafficked, leading to accumulation in the cytoplasm and promoting formation of ubiquitinated TDP-43 aggregates (**Figure 5B**, right).

**Figure 5.**
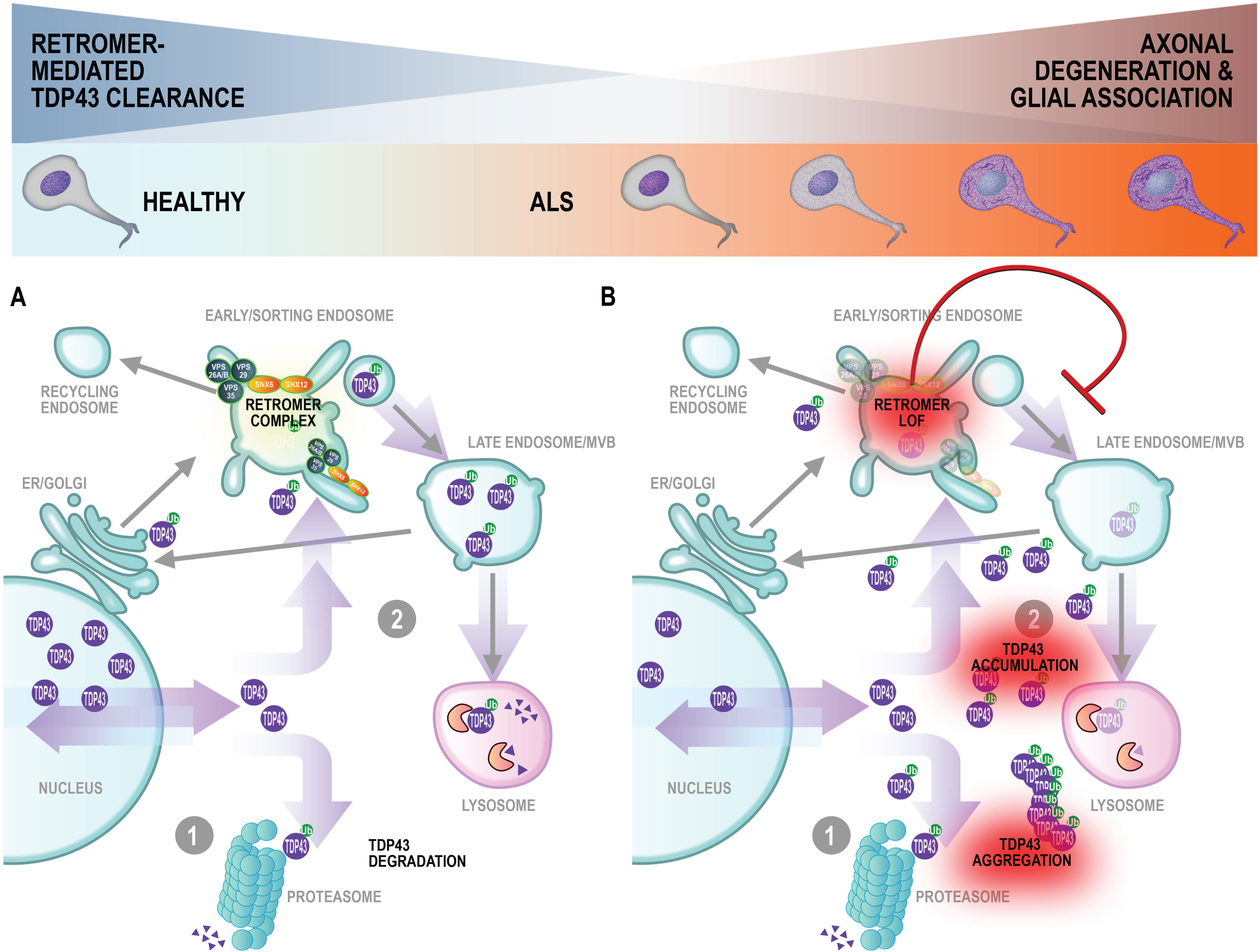
Modeling motor neuron degeneration in TDP-43 pseudotime: early and sustained retromer complex dysfunction promotes accumulation of TDP-43 accompanied by modulation of the neuronal-glial axis. Maintenance of proteostasis in **(A)** healthy MNs through clearance of cytoplasmic TDP-43 via (1) proteosomal degradation or (2) recycling through the endo-lysosomal pathway. Protein sorting is carried out in the early endosome via activity of cargo selective vacuolar sorting proteins (VPS) and nexins (SNX) for retrograde trafficking or lysosome-mediated degradation **(B)** Diminished abundance of VPS and SNX proteins in early ALS (NON) leads to inefficient protein sorting and trafficking, promoting cytoplasmic accumulation and aggregation of TDP-43 proteins\

### Stress granule protein abundances correlate inversely with cytoplasmic TDP-43 inclusions

We observed abundance reductions in stress granule and aggresome proteins G3BP1, USP10, and ATXN2 that inversely correlate with TDP-43 cytoplasmic inclusions. Consistent with the decreased abundance of USP10 and ATXN2 in ALS MNs (**Figure 4C*(ii)***), the ubiquitin-specific protease USP10 has previously been shown to promote clearance of TDP-43-positive stress granules in cells treated with proteasome inhibitor and promote formation of TDP-43-positive aggresomes, while depletion of USP10 increased cytoplasmic insoluble TDP-43 cleavage products (Takahashi et al., 2022). ATXN2 (Ataxin 2) protein has been shown to be diminished in abundance and to colocalize with phosphorylated TDP-43 in hippocampal neuronal cytoplasmic inclusions (NCIs) and dystrophic neurites in FTLD-TDP brains (Watanabe et al., 2020), while intermediate-length repeat expansions in ATXN2 are associated with increased ALS risk (Elden et al., 2010; Lee et al., 2011).

### Neuronal and astrocyte proteins increased with TDP-43 inclusion abundance

A subset of 15 proteins with increased abundances in ALS MN samples demonstrated significant positive correlations with the severity of cytoplasmic TDP-43 inclusions **(Figure 4E),** including 7 glial proteins. Glial:neuronal enrichment scores that we calculated based on cell type-specific proteomic measurements from mouse brains (Sharma et al., 2015) suggest that these proteins, including GFAP and SYNM, are consistent with astrocyte contamination of our MN captures (Krach et al., 2018) and may be indicative of increased astrocyte activity/reactivity in the MN neurodegenerative process (**Figure 4E-F**). CNTNAP2 (contactin-associated protein 2, CASPR2) was the most neuron-enriched protein to show a positive correlation with increasing presence of cytoplasmic TDP-43 inclusions. CNTNAP2 is a transmembrane neurexin protein localized in potassium channel-rich axonal juxtaparanodes with important roles in mediating cell:cell interactions (Poliak et al., 1999; Poliak et al., 2001) and regulation of axonal excitability (Scott et al., 2019). Notably, peptides from the CNTNAP2 extracellular domain have also been detected in human CSF (Martin-de-Saavedra et al., 2022), including in CSF from ALS and FTLD patients (Oh et al., 2021 (preprint)).

### Comparison to ALS proteomic and transcriptomic studies

Mitochondrial and translation impairment in ALS, suggested by the reductions in protein expression profoundly affecting oxidative phosphorylation and ribosome complexes, have been previously implicated by proteomic studies of bulk spinal cord and brain tissues from ALS/FTLD donors (Engelen-Lee et al., 2017; Hartmann et al., 2018; Iridoy et al., 2018; Ladd et al., 2017; Umoh et al., 2018). Reminiscent of these findings, Tank and colleagues described reduced stability of transcripts encoding many oxidative phosphorylation and ribosomal proteins in ALS patient-derived fibroblasts, iPSCs and brain/spinal cord tissues (Tank et al., 2018). A transcriptomic dataset from human FTLD cortical cells identified differentially expressed transcripts associated with loss of TDP-43 nuclear immunoreactivity, a subset of which were identified as splice targets of TDP-43(Liu et al., 2019; Ma et al., 2022). We identified peptides corresponding to 16 of 66 transcripts that undergo alternative splicing with loss of TDP-43 nuclear immunoreactivity, 12 (75%) of which showed altered protein abundances (11 reduced: CADPS, CAMK2B, CYFIP2, EIF4G3, EPB41L1, HP1BP3, IMMT, KIF3A, PRUNE2, STXBP1, SYNE1, UQCRC2; 1 increased: ARHGEF11) prior to detection of cytoplasmic TDP-43 inclusions in MNs in our study. The utility of correlating RNA to protein abundance measurements has been the topic of significant interest and debate in the field (Edfors et al., 2016; Lundberg et al., 2010; Maier et al., 2009; Payne, 2015; Petryszak et al., 2016), ultimately suggesting their complementarity and that RNA:protein data must be interpreted on a case by case basis to shed light on the mechanistic regulation of a given locus. Interestingly, transcriptomic studies of LCM lumbar MNs from patients with bulbar-onset ALS, which could represent an less-degenerated “earlier” disease state than the MNs examined in our study, reported widespread increased abundance of mRNAs in ALS MNs (Krach et al., 2018; Rabin et al., 2010).

### Limitations of this Study

As a nascent field, we should expect single cell proteomic approaches to render greater understanding of disease biology as the technologies mature. In the first known application of single cell proteomics to MNs in ALS, we encountered several challenges. Notably, MN proteome coverage was low at ∼2500 proteins, estimated at about 20-25% coverage of the human core proteome (Wilhelm et al., 2014). Although we measured clear, consistent, and biologically relevant differences in abundances of about 500 proteins between control and ALS MNs, these challenges clearly limited the discovery of low abundance proteins that may be altered in the neurodegenerative process. Furthermore, due to the low throughput capacity of our methods, we analyzed only 135 MNs from 6 ALS donors in our two proteomics experiments. While we increased our per group sample size significantly in our TDP-43-focused study relative to our pilot based on measured (**Supplemental Figure S3**) and modeled (Boekweg et al., 2021) protein variability measured across control cells, we found that our sample size was still too small for comparisons across groups of MNs with more prominent cytoplasmic TDP-43 inclusions. It is not difficult to imagine that the widespread alterations in protein abundances early in the neurodegenerative cascade could exacerbate proteome variability in subsequent disease stages, even among cells that look similar with respect to histologically-detected TDP-43.

Autopsy tissues provide a cross-sectional sample of cells at an advanced stage of human disease. Like other deep molecular phenotyping technologies, single cell proteomic trajectory analysis may soon provide the capability to delineate molecular stages of disease progression without true longitudinal tissue sampling. We employed immunohistochemical assays in adjacent frozen sections to stratify MNs across four phenotypic stages based on the presence of cytoplasmic TDP-43 inclusions, operating under the assumption that increasing presence of histologically-evident TDP-43 inclusions indicates more advanced stages of neurodegeneration. With our approach and sampling depth, we were unable to delineate cell trajectories *de novo* using the CellTrails (Ellwanger et al., 2018) algorithm for single cell trajectory analysis that has been employed in studies of cell differentiation (data not shown), which may be related to the small number of single cells and relatively few associated features in our single cell proteomics dataset compared to single cell RNAseq studies (Chen et al., 2019). Alternatively, we identified altered-in-abundance proteins with the highest Spearman correlations with TDP-43 strata as those that may play important mechanistic roles in ALS. However, our assumption of the direct relationship between a MN’s neurodegenerative progression and observable cytoplasmic TDP-43 inclusions density does not take into account the effects of oligomer toxicity or the potential neuroprotective roles of large aggregates (Arrasate et al., 2004; Subramaniam et al., 2009).

A further limitation of our study stems from its reliance on banked human autopsy specimens, which imparts additional variability in sample handling outside of our control. Autopsy and tissue banking protocols vary from site to site, and while we ensured that ALS and CTL tissues in our study originated from the same site, we nonetheless observe significant variation in the postmortem interval prior to sample preservation (**Figure 1A**). While we tried to match samples based on available demographic information, we were limited by sample availability in our ability to perfectly match our CTL and ALS donors with respect to potential covariates (**Supplemental Table S8**). To explore the effects of these potential confounders on identification of differentially abundant proteins, we performed parallel analyses of our ALS Pilot dataset with and without correction for age, sex, and PMI covariates using a linear mixed model (*limma*) (Ritchie et al., 2015) (**Supplemental Figure S4, Supplemental Figure S5**). We observed clear overlap in the identifies of differentially abundant proteins determined using limma and by T-test (**Supplemental Table S1**). Notably, however, we observed a striking reduction in the number of proteins meeting the threshold for statistically significant differential abundance (**Supplemental Figure S4B**). While only 7 proteins survived this stringent correction – SNRPD1, CYCS, EEF1A1, H2AZ2, FUBP1, and CAPRIN1 –these proteins confer provide additional support for pathway-level dysregulation of cellular respiration and oxidative phosphorylation, RNA splicing, and protein translation in ALS MNs (**Supplemental Figure S4C**).

## Conclusion

Herein we have presented, to our knowledge, the first unbiased proteomic studies of single somatic MNs from postmortem ALS donor spinal cords. Trace-sample protein processing using NanoPOTS: (1) rendered identification of approximately 2500 proteins and quantitative comparison of approximately 1000 proteins in our pilot study; (2) enabled disease-state differentiation of single MNs; and (3) revealed reductions in protein abundances suggestive of impairments in cell energetics, protein translation, proteostasis and trafficking mechanisms, including a new emphasis on Golgi-lysosome trafficking. In a follow-up experiment in which we stratified MNs by TDP-43 neuropathology, we observed similar patterns of differentially expressed proteins in MNs lacking TDP-43 NCIs, suggesting that impairments in cellular energetic and proteostastic mechanisms occur early with respect to formation of TDP-43 inclusions. Specifically, we report declining expression of retromer complex components accompanying increasing presence of TDP-43 inclusions and propose retromer complex-mediated endolysosomal sorting as a potential point of future mechanistic research in the formation or clearance of TDP-43 inclusions in vulnerable ALS motor neurons. We further confirmed loss of canonical STMN2 transcripts and protein in ALS MNs concurrent with early TDP-43 pathology, indicative of loss of TDP-43-mediated splicing functions prior to formation of large NCIs. Our study begins to show the potential of single cell proteomics to augment the study of human neurologic diseases and provide insights into temporal dynamics of disease progression directly from human tissues.

**Table 1.** Patient Characteristics. Demographic data and characteristics of sampled patients. Statistical comparisons were determined using student’s t-test for numerical values or chi-squared test for categorical values.

|  | ALS<br>(N=8) | CTL<br>(N=6) | Overall<br>(N=14) | P-Value |
| --- | --- | --- | --- | --- |
| **PMI..hours.** |  |  |  |  |
| Mean (SD) | 15.3 (10.2) | 15.7 (3.27) | 15.5 (7.79) | 0.937 |
| Median [Min, Max] | 10.7 [5.00, 30.7] | 14.5 [12.9, 21.0] | 13.5 [5.00, 30.7] |  |
| **Age..years.** |  |  |  |  |
| Mean (SD) | 71.3 (8.01) | 55.5 (6.38) | 64.5 (10.8) | 0.00192 |
| Median [Min, Max] | 71.0 [59.0, 81.0] | 57.0 [46.0, 63.0] | 63.5 [46.0, 81.0] |  |
| **Sex** |  |  |  |  |
| F | 4 (50.0%) | 1 (16.7%) | 5 (35.7%) | 0.469 |
| M | 4 (50.0%) | 5 (83.3%) | 9 (64.3%) |  |

## Acknowledgements

We express our gratitude first and foremost to the individuals and families who have generously donated their tissues for the advancement of medical research, and to the University of Miami Brain Endowment Bank™, which is funded by NIMH, NINDS, and NICHD, for collection and provision of samples. We additionally thank members of the Biogen Neuromuscular Disease Research Unit, Translational Biology, Biostatistics, Translational Neuropathology, and Biomarker teams for their insightful discussions and feedback. This research was supported by National Cancer Institute of the National Institutes of Health awards R33 CA225248 and R01 GM138931 to RTK and through a sponsored research agreement with Biogen.

## Author Contributions

RTK, EDP, AJG, and SAM conceived and planned the experiments. SAM, RC, TT, YL, RTK, AJG, and EDP collected the data. AJG, SAM, RC, HB, JWC, NCW, JG, DVDW, SHP, RTK, and EDP analyzed and interpreted the results. AJG, SAM, RC, RTK, and EDP wrote the manuscript with critical input and feedback from all authors.

## Supplemental Figure Legends

**Supplemental Figure S1.**
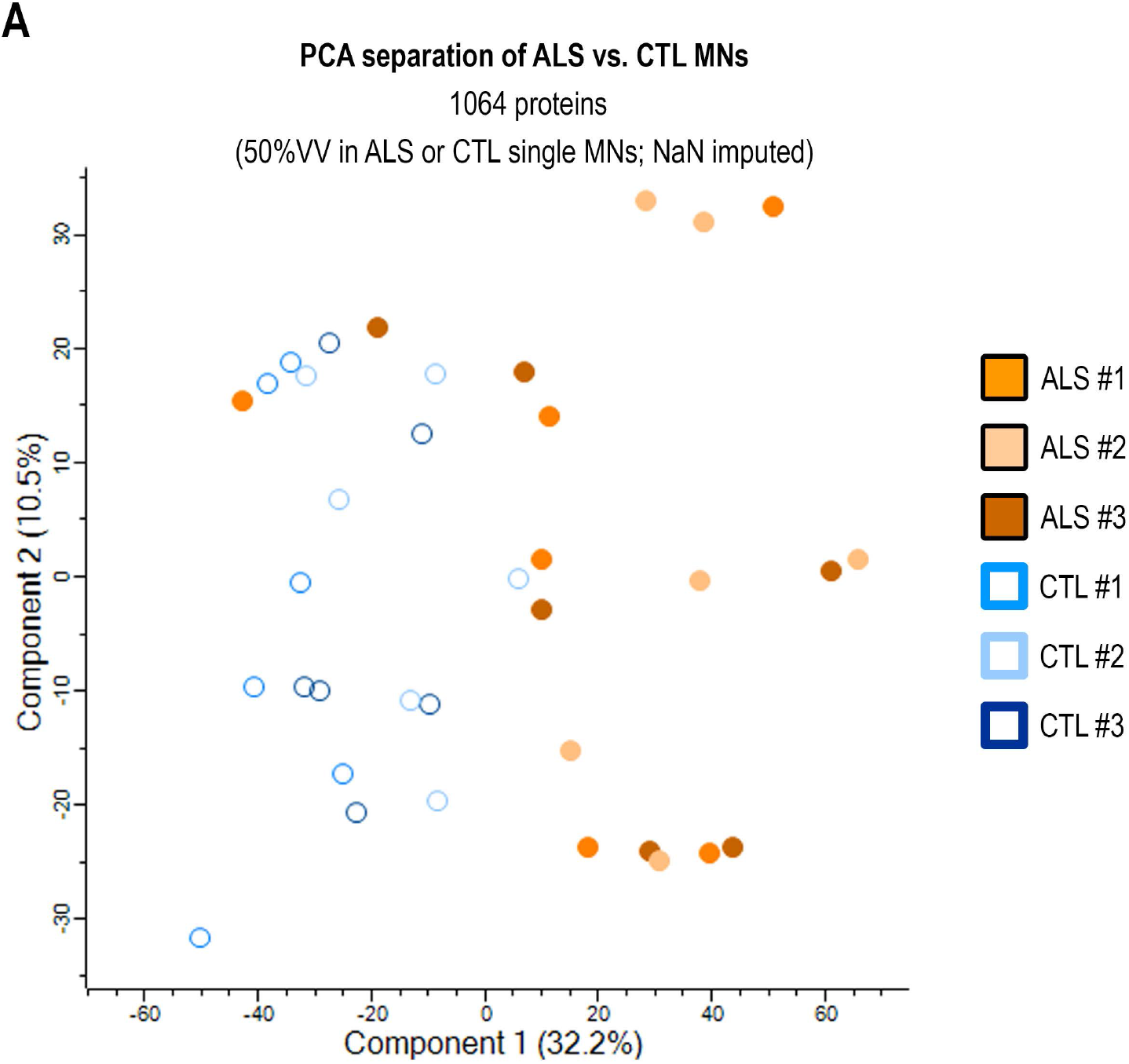
Separation of ALS and CTL MNs based on single cell proteome measurements. Dimensionality reduction by principal component analysis of 1064 protein abundances measured across ALS and CTL MNs from n=3 donors per group (ALS/CTL #1-#3).

**Supplemental Figure S2.**
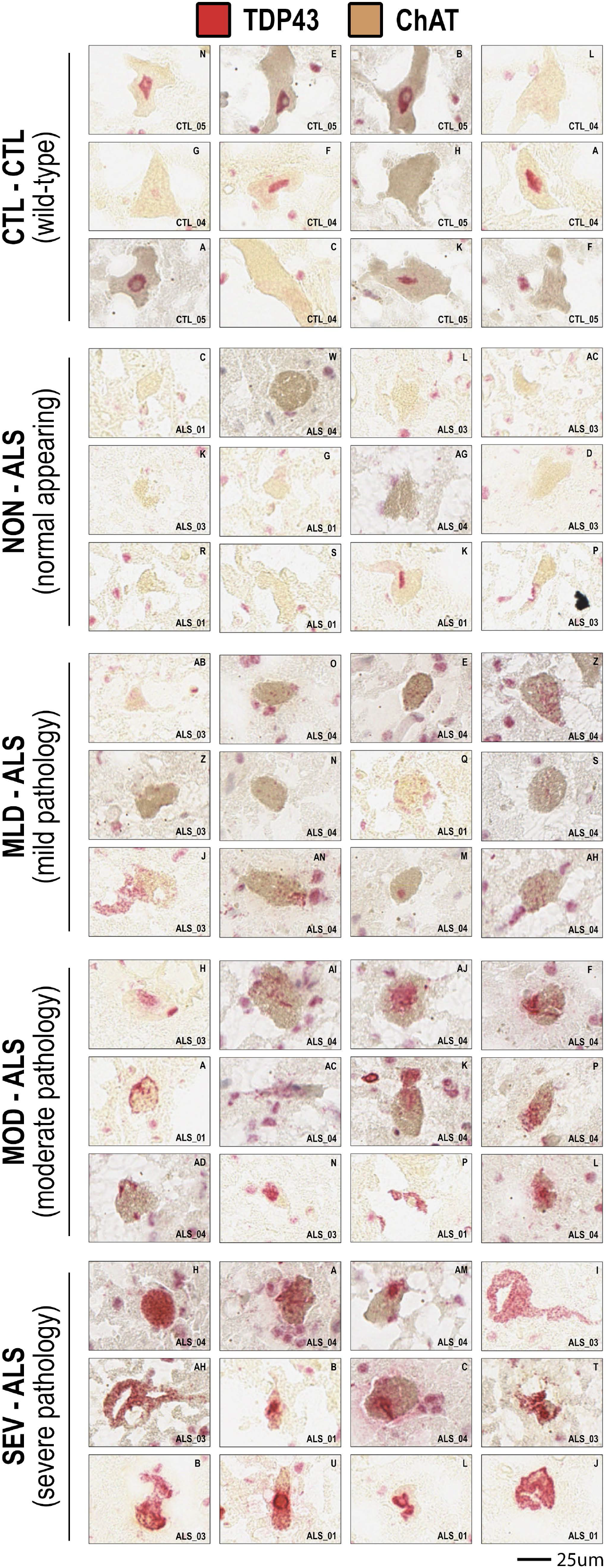
TDP-43 histopathology staging across ALS motor neurons. **(A)** Representative TDP-43 immunohistochemistry staining for individual MNs selected for nanoPOTS LC-MS/MS analysis. TDP-43 Inclustion strata Stages : CTL, CTL – normal appearing from healthy control donors; NON-ALS – normal appearing from ALS donors; MLD-ALS – mild inclusions/pathology from ALS donors; MOD-ALS – moderate inclusions/pathology from ALS donors; SEV-ALS – severe inclusions/pathology from ALS donors **(B)** Distribution of MNs per donor across defined TDP43 strata

**Supplemental Figure S3.**
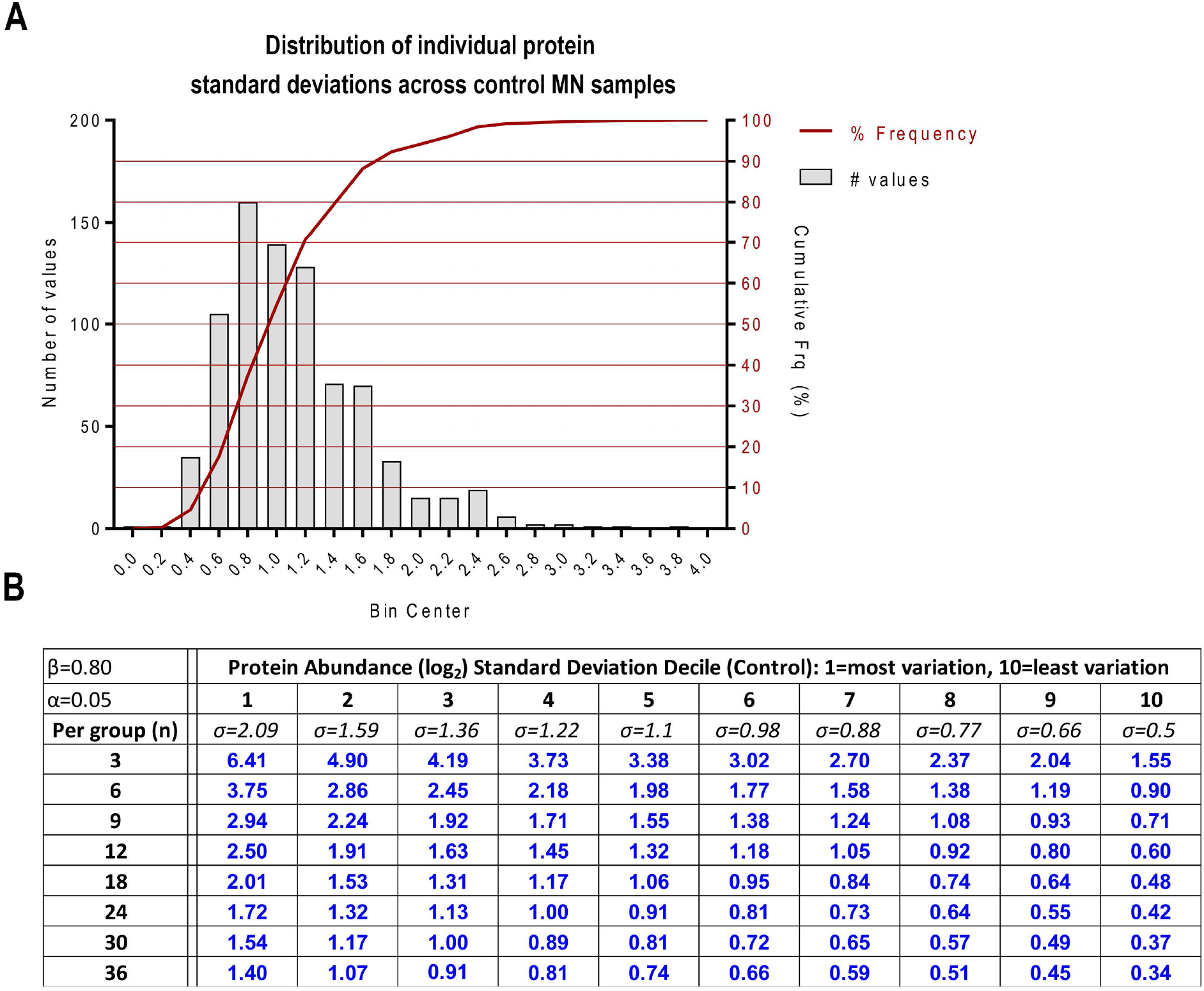
Determination of protein abundance variability across individual healthy motor neurons. **(A)** Distribution of standard deviations per protein measured across healthy CTL MNs samples (n=18) **(B)** Predicted range in protein variability measurements by standard deviation decile based on measurements from CTL MNs samples for multiple samples sizes (Type I error = 0.05; Power = 80%).

**Supplemental Figure S4.**
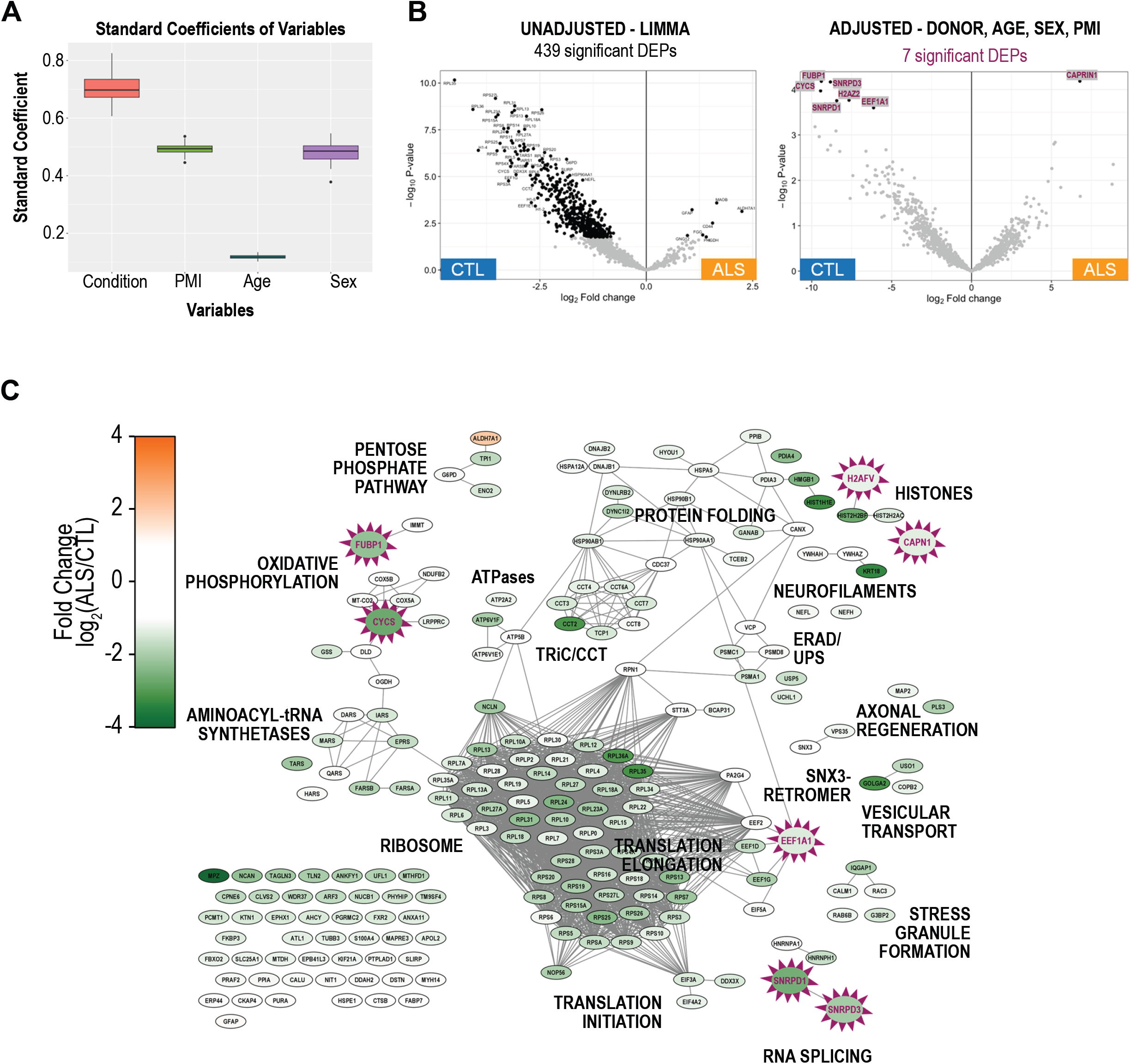
Effect of donor demographic variables on significance determinations for differential protein abundance comparisons in ALS and CTL MNs. **(A)** Standard coefficient of variables (condition - ALS or control postmortem interval (PMI), age, and sex) associated with MN donors **(B)** Comparison of differential protein abundance determinations with or without adjustment for donor variables using linear mixed models (limma) **(C)** Overlay of proteins surviving correction for donor variables (n=7, highlighted nodes) on the background of differentially abundant proteins deemed significance without correction for donor variables.

**Supplemental Figure S5.**
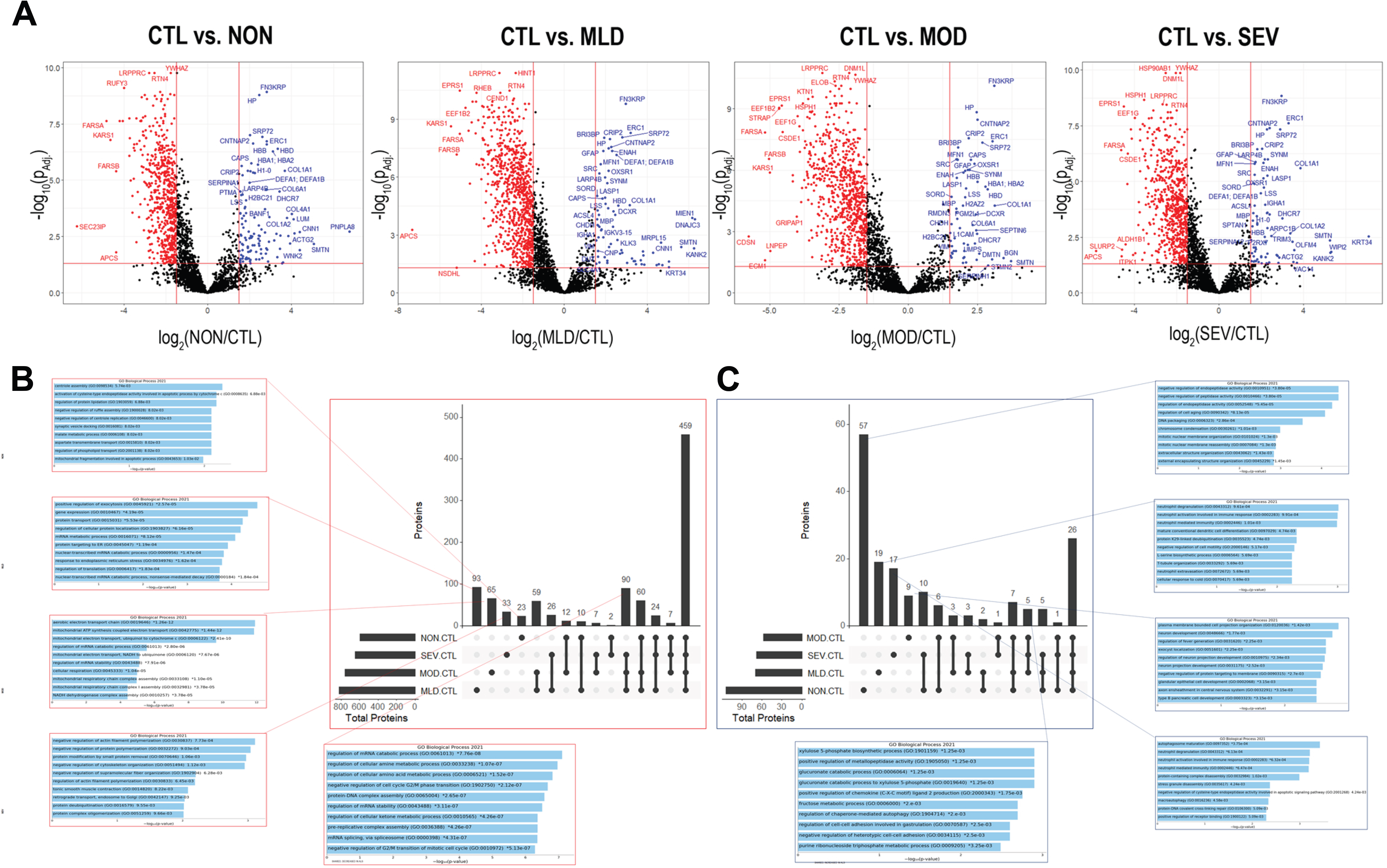
Effect of donor demographic variables on significance determinations for differential protein abundance comparisons across TDP43 strata. **(A)** Volcano plots of differential protein abundance comparisons for individual TDP43 strata (mild – MLD, moderate – MOD, severe – SEV) compared to control MN (NON); Comparisons were carried out using linear modeling (fold change cutoff: Log_2_FC>1.5, Benjamini-Hochberg-adjusted p-value <0.05) **(B-C)** Upset plots highlighting shared and uniquely differentially abundant proteins across TDP43 strata for proteins (B) decreased in abundance in ALS MNs and **(C)** increased in abundance in ALS MNs with corresponding top-10 over-represented Pathological GO terms as identified using EnrichR (bar graphs) and corresponding adjusted p-values.

## Supplemental Table Descriptions

*Supplemental Table S1: High confidence master proteins identified in human ALS or CTL MNs (TDP-43-agnostic)*

*Supplemental Table S2: Significantly differentially abundant proteins identified in human ALS vs. CTL MNs (TDP-43-agnostic)*

*Supplemental Table S3: High confidence master proteins identified in human ALS MNs stratified by TDP-43 histopathology*

*Supplemental Table S4: Mean stage-to-stage TDP-43 histopathology stratified abundance values per protein in ALS MNs*

*Supplemental Table S5: Stage-to-stage TDP-43 histopathology stratified protein abundance fold change in ALS MNs*

*Supplemental Table S6: “Early” and “late” subsets of differentially abundant proteins in ALS MNs*

*Supplemental Table S7: Individual protein abundance correlations (Spearman) with TDP-43 histopathology stage*

*Supplemental Table S8: Patient characteristics for ALS and CTL human tissue donor samples*

## Methods

### Case selection

ALS and CTL cases were selected from a cohort of frozen thoraco-lumbar spinal cord tissue samples from the University of Miami Brain Bank (**Supplemental Table S8**). Spinal cord tissues were divided into ∼3mm sub-blocks. Two to seven sub-blocks were generated per sample, depending on initial tissue dimensions. Sub-blocks were generated by allowing the tissue to thaw slightly to avoid fracturing, then sliced into sub-blocks using a razor blade. Sub-blocks were immediately transferred to histology cassettes on dry ice and stored at −80 °C. One ∼3mm sub-block per donor was embedded in pre-chilled 2.5% carboxymethylcellulose solution (2.5% (w/v) carboxymethylcellulose (Sigma, Cat. No. 419273-100G, Lot. No. MKCG5725) in dH_2_O) in cryomolds on dry ice and allowed to freeze fully at −80 °C. Subsequently, 5um sections were collected on positively-charged histology slides (Fisher Superfrost) and post-fixed by immersion in 10% Neutral Buffered Formalin (Fisher Scientific) for 15 minutes followed by two 10 minute washes in 1X phosphate buffered saline (PBS) (Abcam, Ref. #ab128983). All sections were collected using a Leica CM1520 cryostat (CMMS ID XX-99339) using a Leica high-profile microtome blade at an internal cryostat temperature of −23C. Fixed slides were allowed to air dry following PBS rinse and stored at room temperature prior to staining. For tissue evaluation and case selection, hematoxylin and eosin (H&E) staining was carried out a single frozen section per donor using an automated Leica Spectra staining protocol (H&E no oven) and coverslipped using an automated coverslipper (TissueTek). Brightfield images of stained tissues were collected of all tissues at 20× on 3DHISTECH Panoramic slide scanner. In addition to evaluation of tissue quality, sample were selected to minimize differences in age at time of autopsy and postmortem interval (PMI) across sample groups.

### Tissue sectioning and laser capture microdissection

Zeiss PEN membrane slides (Zeiss, Ref # 415190-9041-000, Lot # 000671-19) and Superfrost PlusGold slides (Fisher, Ref # 15-188-48, Lot # 19906-665158) were exposed to UV light for 30 min in a laminar flow hood to promote tissue adhesion. Spinal cord blocks were transferred from the −80 °C freezer to the cryostat (−23 °C) and allowed to equilibrate in temperature for 20 min. 12-µm-thick sections were collected by cryosectioning in the following order: (1) one 12 µm section on a Superfrost PlusGold slide; (2) ten adjacent 12 µm sections (two per slide) on five PEN membrane slides; (3) one 12 µm section on a Superfrost PlusGold slide.

Slides were pre-chilled on the interior surface of the cryostat prior to section collection. Single sections were picked up on PlusGold slides, adhered to the slide by warming the reverse side of the slide with a finger, and then immediately refrozen on the inner cryostat surface. Two sections were collected at the same time onto each PEN membrane slide, melted onto the slide by warming the reverse side of the slide with a finger, and then were immediately refrozen on the inner cryostat surface. Once sections were frozen and all sections (A-C) for a sample were collected, tissue sections were fixed in 70% ethanol for 15 minutes at room temperature and then transferred immediately to storage at −80 °C in slide boxes with desiccant packs (Humidity Sponge).

### For TDP-43-agnostic MN captures (ALS Pilot)

all sections were collected over the course of four days, ensuring that equal numbers of ALS and non-disease control cases were collected in individual sectioning sessions using the same cryostat. All sections were stored at −80 °C prior to H&E staining, ethanol dehydration, and vacuum desiccation. Stained tissue sections were scanned on a Zeiss PALM MicroBeam system at 40× resolution. Motor neurons were selected for single cell based on morphology and presence of Nissl substance and if the same cell was confidently identified across two or more adjacent sections to allow capture of two 12um-thick excised cell cross-sections per sample. Six individual MNs from each case (3 ALS and 3 CTL) were selected for subsequent analysis. Specifically, individual MNs matched across two adjacent sections were selected from laminae VII/IX of the ventral horn, excised by laser capture microdissection (LCM), and collected in individual nanoPOTS wells prefilled with DMSO to aid in sample collection using the Slide Collector 48 adapter. 10MN boost samples were generated for each donor by capturing and combining twenty 12um-thick excised cell cross-sections to approximate the protein contents of ten cells (two cross-sections per cell). Nanowells were imaged at 10× resolution to confirm collection of each excised cell. Following collection, nanoPOTS chips were sealed to avoid the evaporation of the solution from the nanowells.

### For TDP-43-pathology-guided MN captures (TDP-43 Pseudotime)

20um-thick frozen sections were collected onto Zeiss PEN membrane slides, as described above, and adjacent 10um-thick frozen sections were collected onto Plus Gold slides for subsequent chromogenic detection of TDP-43 and ChAT protein by immunohistochemistry (IHC) staining. Frozen human spinal cord cross sections were fixed in 10% NBF, dried, and stored at room temperature prior to immunostaining. Staining was conducted on a Ventana Ultra platform using standard chromogenic methods. For antigen retrieval (HIER) and permeabilization, slides were heated in a pH9 EDTA-based buffer for 10m at 94°C, followed by incubation with a mouse monoclonal antibody against TDP-43 (L95A-42, Biogen) at 1:40,000 and a rabbit monoclonal antibody targeting ChAT (Clone: EPR16590, Abcam, Ref. ab178850) at 1:3,000. Bound anti-TDP-43 and anti-ChAT primary antibodies were detected using an AP-conjugated-anti-mouse and HRP-conjugated-anti-rabbit secondary polymers with chromogenic visualization with Ventana DISCOVERY Red and Ventana DISCOVERY Yellow, respectively. A subset of slides were counterstained with hematoxylin to visualize nuclei. Stained slides were imaged at 20×. Images from adjacent H&E-stained 20um- and TDP-43-ChAT-stained 10um-sections were manually aligned for identification of single MNs spanning both sections for laser capture microdissection (as described above) and classification of TDP-43 pathology, respectively. TDP43 strata were defined along a semi-quantitative 4-point scale based on quantity and morphology of cytoplasmic TDP43 protein inclusions:

### NanoPOTS sample processing

Following cell capture, remaining DMSO was allowed to evaporate prior to adding additional processing reagents. Samples were further processed using the nanoPOTS workflow as described previously (Cong et al., 2020a; Zhu et al., 2018a; Zhu et al., 2018c). Briefly, proteins were extracted with 0.1% dodecyl-β-d-maltopyranoside (DDM) and reduced with 5 mM dithiothreitol (DTT) followed by alkylation with 10 mM iodoacetamide (IAA). The two-step enzyme digestion was performed with LysC (0.25 ng) for 4 h followed by trypsin (0.25 ng) for an additional 16 h at 37 °C. Digestion reactions were quenched with 0.1% formic acid (FA) and digested peptide samples were collected in 200-µm-i.d. fused silica capillaries using a robotic liquid handling system. The samples were stored individually at −20 °C prior to LC-MS/MS analysis. Additional “boost” samples (10 MN equivalents) were prepared using pooled MNs from each case to facilitate feature identification and matching across single-cell analyses of the same case (Figure S1). Prior to injection and MS analysis, samples were block-randomized to minimize batch effects and impact of instrument drift during data acquisition.

### nanoLC-MS/MS analysis

Samples were equilibrated to 4°C prior to analysis and were positioned for in-line loading on to an in-house-packed SPE column (5 cm x 75-µm-i.d.). Samples were loaded onto the column over 10 min using 100% Mobile Phase A (0.1% FA in water) at a flow rate of 0.5 µL/min using an UltiMate 3000 RSLCnano pump (Thermo Fisher) to ensure complete desalting. Peptide separation was carried out by connecting the SPE column to an in-house-packed analytical SPE column (50 cm x 30-µm-i.d.) connected to a nanospray emitter by a zero-dead-volume union (Valco, Houston, TX). Peptides were separated by a 100-min linear gradient (8-25% mobile phase B (0.1% FA in acetonitrile) followed by an additional 20-min linear gradient (25-45% B) to elute hydrophobic peptides. For column washing, mobile phase B was increased to 90% over 5 min and held constant for 5 min to wash the column, then was reduced to 2% over 5 min and held constant for 15 min to re-equilibrate the column. Post-split (50 cm x 75-µm-i.d. split column) mobile phase flow rates were 20 nL/min; 250 nL/min programmed flow was provided by the UltiMate 3000 RSLCnano pump.

Peptides were injected into a Thermo Orbitrap Exploris 480 mass spectrometer by electrospray using in-house-pulled nanospray emitters (20-µm-i.d.). MS and MS/MS data were acquired by employing an electrospray potential of 2000 V at the source for ionization. The ion transfer tube temperature was 200 °C for desolvation and the ion funnel RF level was 40. Full MS scans were acquired at 375-1800 m/z with an orbitrap resolution of 120,000 (*m/z* 200). The AGC target and maximum injection time were set to 1E6/200 ms. Data-dependent MS/MS spectrum acquisitions were triggered for precursor ions with intensities > 5E3 and charge states of +2 to +7. The scan range was defined from first mass to 100 m/z with a cycle time of 3 sec. Monoisotopic precursor ion peaks were fragmented by higher energy collision-induced dissociation (HCD) with a normalized collision energy of 28% and with AGC target and maximum injection time set to 1E5/500 ms. Fragment ions were detected in the orbitrap at 30,000 resolution (*m/z* 200). MS/MS isolation windowswere 1.6 Da with a mass tolerance of ±10 ppm and dynamic exclusion time was set to 90s.

### Protein identification and quantitation

Raw MS data were searched against a protein database consisting of reviewed human proteins (20,353 reviewed protein sequences, UniProtKB, downloaded: July 20^th^, 2020) appended with common contaminants using a two-step database search was done with Sequest HT and Sequest HT INFERYS rescoring algorithms in Proteome Discoverer (version 2.5, ThermoFisher Scientific, San Jose, CA) specifying fully tryptic enzymatic digestion (7-30 amino acids, 2 missed cleavages). Fixed carbamidomethylation (C) and variable oxidation (M), deamidation (N,Q) and modification of protein N-termini (acetyl, Met-loss, pGlu) were included in the search parameters as modifications. Precursor and fragment mass tolerances were set to 10 ppm and 0.02 Da, respectively. The peak matching feature detection option was enabled to allow a maximum chromatographic retention time shift to 10 min with a mass tolerance of 10 ppm. Peptide identifications were refined using a target-decoy approach followed by percolation based on q-values and imposing a strict FDR cut-off of 0.01 and a relaxed FDR cut-off of 0.05 at the PSM and peptide level. Protein abundances were determined using the precursor ions quantifier node based on the top 3 distinct peptides (unique and razor) from each protein and normalized to total peptide signal per single cell sample (“boost” samples were included in the search to facilitate protein identification via feature matching but were excluded in normalization and quantitative comparison steps).

### Data analysis and visualization

Protein differential expression analysis between ALS and CTL MNs was performed using scripts in R (version 4.1.0) (R Core Team, 2020) as follows: a paired t-test using the t.test() function from the stats package was used to determine p-values for fold-change protein abundance differences between ALS and CTL MN data; p-values were then adjusted for multiple comparisons using the *p.adjust()* function (Benjamini-Hochberg/FDR option). Proteins with an adjusted p-value < 0.05 and a fold-change >2 or <0.5 were designated as differentially expressed. Depiction of the differential expression analysis as volcano plots was performed using the *ggplot2* (Wickham, 2016) package in R. Fold-change quantification data for proteins identified as differentially expressed between ALS MN and CTL neurons was grouped by TDP-43 pathology stage for protein trajectory analyses. 95% confidence intervals for the fold-change values were computed according to Fieller’s Method.

Interaction networks for proteins exhibiting significantly differential abundance in ALS vs. CTL MNs were generated in the web-based STRING application (v11.5)(Szklarczyk et al., 2019) with additional filtering requiring a minimum interaction score of 0.7 and with “Experimental” and “Database” active interaction sources enabled. Networks were then imported into Cytoscape (v3.8.2)(Shannon et al., 2003) to allow mapping of protein abundance data onto individual nodes. For TDP-43 negative correlators, the physical interaction network was constructed as described above, with the following modifications: minimum interaction score = 0.4; “physical subnetwork” option enabled. Over-representation of gene ontology, KEGG, and Reactome pathway terms associated with identified protein subsets were determined using hypergeometric tests (statistical background = whole genome) followed by Benjamini-Hochberg correction for multiple hypothesis testing using the STRING enrichment analysis widget (Franceschini et al., 2013). Pearson and Spearman correlation scores were calculated using the PerseusGUI (v1.6.5.0)(Tyanova et al., 2016). Glial:neuronal enrichment scores were calculated based on published data from a study focused on comprehensive profiling of the mouse brain proteome across cell types and brain regions. Median abundance values were calculated across all ages/DIV stages for each cell population: astrocyte, oligodendroglial, microglial, and neuronal LFQ (label-free quantitation) data published in *Sharma, et al. 2016, Table S1* (File ID: https://static-content.springer.com/esm/art%3A10.1038%2Fnn.4160/MediaObjects/41593_2015_BFnn4160_MOESM38_ESM.xlsx; <u>PXD001250</u>; (Sharma et al., 2015)). Enrichment scores per population were then calculated as the log_2_-transformed ratios of median abundance in each glial population vs. the median abundance in the neuronal population. A pan-glial ratio for each protein was also calculated as the mean of the individual per-glial-subtype enrichment scores. Resulting data were analyzed using GraphPad Prism.

For TDP43 stratified samples, correction for individual donor contribution was evaluated by considering donor origin (n=3) of each ALS MN sample as individual variables and control donor origin (n=2) as a single variable. Raw protein intensity data were log_2_-normalized and compared across each stratified TDP43 group (i.e., NON, MLD, MOD, SEV) to the control and to each other, accounting for donor contribution as a variable. Differential protein abundance between groups was determined using LIMMA and p-values were corrected for multiple hypothesis testing (Benjamini-Hochberg). Significantly differentially abundant proteins (adjusted p-value <0.05 and logFC >1.5) were quantified for each comparison and visualized using volcano plots in R. Lists of significantly increased- or decreased-in-abundance proteins were made for each TDP43 stratum relative to the control group. Using these lists, we compared the unique proteins (only significant in one TDP43 stratum) via upset plot separately for each differentially abundant protein subset. Unique proteins were subjected to pathological ontology analysis, weighted by logFC using EnrichR (https://maayanlab.cloud/Enrichr/) and top 10 terms with lowest p values were displayed. All statistical analyses were done in R.

### In-situ hybridization and immunohistochemistry staining

FFPE tissue sections from human spinal cord samples were evaluated for RNA quality using positive (BA-Hs-POLR2A-3zz) and negative control (BA-dapB-3zz) BaseScope (Advanced Cellular Diagnostics, Inc.) probes. Expression of canonical STMN2 transcripts was investigated using a custom (BA-Hs-STMN2-3zz-st) 3zz BaseScope probe (Advanced Cellular Diagnostics, Inc.) in conjunction with detection of TDP-43 protein using a mouse antibody against TDP-43 (0.25 ug/ml, TDP-43-L95A, Biogen) and Bond Primary Antibody Diluent (Advanced Cellular Diagnostics, Inc., Cat. No. AR9352) and the Bond Polymer Refine Detection Kit (Advanced Cellular Diagnostics, Inc., Cat No. DS9800).

RNA *in situ* hybridizations were performed on a Leica Bond automated platform using the BaseScope Reagent Kits (Advanced Cell Diagnostics, Inc.) according to manufacturer’s instructions. Briefly, 5um-thick FFPE tissue sections were pretreated with heat (Epitope retrieval (LS ER2) was carried out for 30 minutes at 95C) and protease (Protease IV, 30 min at 40C) prior to probe hybridization and antibody incubation (15 minutes, 0.25ug/ml). Pre-preamplifier, preamplifier, amplifier, and HRP/AP-labeled oligos were hybridized sequentially, followed by chromogenic detection. Samples were counterstained with hematoxylin. Brightfield images were collected at 40X using a 3DHistech Pannoramic SCAN II digital slide scanner. Scanned images were processed using a custom image analysis algorithms developed in Visiopharm for threshold-based detection of STMN2 and TDP-43 signal. MNs were annotated by training a Random Forest classifier and the resulting ROIs were manually inspected and adjusted prior to analysis of STMN2 and TDP-43 signal. Resulting data were analyzed using GraphPad Prism.

### Reagents

Mayer’s Hematoxylin solution (MHS16-500ml) and eosin Y solution (110116-500ml) were purchased from Sigma-Aldrich (St. Louis, MO). Scott’s bluing reagent (Cat. No. 6697-32) was purchased from Ricca Chemical Company (Arlington, TX). Ethanol was purchased from Decon laboratories Inc. USA. Pierce HeLa Protein Digest Standard, Pierce Formic Acid (FA) (LC-MS grade), Dithiothreitol (DTT), and iodoacetamide (IAA) were purchased from ThermoFisher Scientific (Waltham, MA). CHROMASOLV™ LC-MS water and acetonitrile were products of Honeywell (Charlotte, NC), MS-grade trypsin and Lys-C were from Promega (Madison, WI). All other chemicals and reagents were pu0072chased from Sigma-Aldrich (St. Louis, MO) unless otherwise noted.

### Data availability

All relevant data are available from the authors upon request.

## References

Arai, T., Hasegawa, M., Akiyama, H., Ikeda, K., Nonaka, T., Mori, H., Mann, D., Tsuchiya, K., Yoshida, M., Hashizume, Y., et al. (2006). TDP-43 is a component of ubiquitin-positive tau-negative inclusions in frontotemporal lobar degeneration and amyotrophic lateral sclerosis. Biochem Biophys Res Commun 351, 602–611.

Arrasate, M., Mitra, S., Schweitzer, E.S., Segal, M.R., and Finkbeiner, S. (2004). Inclusion body formation reduces levels of mutant huntingtin and the risk of neuronal death. Nature 431, 805–810.

Bjork, R.T., Mortimore, N.P., Loganathan, S., and Zarnescu, D.C. (2022). Dysregulation of Translation in TDP-43 Proteinopathies: Deficits in the RNA Supply Chain and Local Protein Production. Front Neurosci 16, 840357.

Boekweg, H., Guise, A.J., Plowey, E.D., Kelly, R.T., and Payne, S.H. (2021). Calculating Sample Size Requirements for Temporal Dynamics in Single-Cell Proteomics. Mol Cell Proteomics 20, 100085.

Chen, H., Albergante, L., Hsu, J.Y., Lareau, C.A., Lo Bosco, G., Guan, J., Zhou, S., Gorban, A.N., Bauer, D.E., Aryee, M.J., et al. (2019). Single-cell trajectories reconstruction, exploration and mapping of omics data with STREAM. Nat Commun 10, 1903.

Cong, Y., Liang, Y., Motamedchaboki, K., Huguet, R., Truong, T., Zhao, R., Shen, Y., Lopez-Ferrer, D., Zhu, Y., and Kelly, R.T. (2020a). Improved Single-Cell Proteome Coverage Using Narrow-Bore Packed NanoLC Columns and Ultrasensitive Mass Spectrometry. Anal Chem 92, 2665–2671.

Cong, Y., Motamedchaboki, K., Misal, S.A., Liang, Y., Guise, A.J., Truong, T., Huguet, R., Plowey, E.D., Zhu, Y., Lopez-Ferrer, D., et al. (2020b). Ultrasensitive single-cell proteomics workflow identifies >1000 protein groups per mammalian cell. Chemical science 12, 1001–1006.

Edfors, F., Danielsson, F., Hallstrom, B.M., Kall, L., Lundberg, E., Ponten, F., Forsstrom, B., and Uhlen, M. (2016). Gene-specific correlation of RNA and protein levels in human cells and tissues. Mol Syst Biol 12, 883.

Elden, A.C., Kim, H.J., Hart, M.P., Chen-Plotkin, A.S., Johnson, B.S., Fang, X., Armakola, M., Geser, F., Greene, R., Lu, M.M., et al. (2010). Ataxin-2 intermediate-length polyglutamine expansions are associated with increased risk for ALS. Nature 466, 1069–1075.

Eleuteri, S., and Albanese, A. (2019). VPS35-Based Approach: A Potential Innovative Treatment in Parkinson’s Disease. Front Neurol 10, 1272.

Ellwanger, D.C., Scheibinger, M., Dumont, R.A., Barr-Gillespie, P.G., and Heller, S. (2018). Transcriptional Dynamics of Hair-Bundle Morphogenesis Revealed with CellTrails. Cell Rep 23, 2901–2914 e2913.

Engelen-Lee, J., Blokhuis, A.M., Spliet, W.G.M., Pasterkamp, R.J., Aronica, E., Demmers, J.A.A., Broekhuizen, R., Nardo, G., Bovenschen, N., and Van Den Berg, L.H. (2017). Proteomic profiling of the spinal cord in ALS: decreased ATP5D levels suggest synaptic dysfunction in ALS pathogenesis. Amyotroph Lateral Scler Frontotemporal Degener 18, 210–220.

Filippone, A., Smith, T., and Pratico, D. (2021). Dysregulation of the Retromer Complex in Brain Endothelial Cells Results in Accumulation of Phosphorylated Tau. J Inflamm Res 14, 7455–7465.

Franceschini, A., Szklarczyk, D., Frankild, S., Kuhn, M., Simonovic, M., Roth, A., Lin, J., Minguez, P., Bork, P., von Mering, C., et al. (2013). STRING v9.1: protein-protein interaction networks, with increased coverage and integration. Nucleic Acids Res 41, D808–815.

Fratta, P., Sivakumar, P., Humphrey, J., Lo, K., Ricketts, T., Oliveira, H., Brito-Armas, J.M., Kalmar, B., Ule, A., Yu, Y., et al. (2018). Mice with endogenous TDP-43 mutations exhibit gain of splicing function and characteristics of amyotrophic lateral sclerosis. EMBO J 37.

Graf, E.R., Heerssen, H.M., Wright, C.M., Davis, G.W., and DiAntonio, A. (2011). Stathmin is required for stability of the Drosophila neuromuscular junction. J Neurosci 31, 15026–15034.

Hartmann, H., Hornburg, D., Czuppa, M., Bader, J., Michaelsen, M., Farny, D., Arzberger, T., Mann, M., Meissner, F., and Edbauer, D. (2018). Proteomics and C9orf72 neuropathology identify ribosomes as poly-GR/PR interactors driving toxicity. Life Sci Alliance 1, e201800070.

Hasegawa, M., Arai, T., Nonaka, T., Kametani, F., Yoshida, M., Hashizume, Y., Beach, T.G., Buratti, E., Baralle, F., Morita, M., et al. (2008). Phosphorylated TDP-43 in frontotemporal lobar degeneration and amyotrophic lateral sclerosis. Annals of neurology 64, 60–70.

Hedl, T.J., San Gil, R., Cheng, F., Rayner, S.L., Davidson, J.M., De Luca, A., Villalva, M.D., Ecroyd, H., Walker, A.K., and Lee, A. (2019). Proteomics Approaches for Biomarker and Drug Target Discovery in ALS and FTD. Front Neurosci 13, 548.

Ho, R., Workman, M.J., Mathkar, P., Wu, K., Kim, K.J., O’Rourke, J.G., Kellogg, M., Montel, V., Banuelos, M.G., Arogundade, O.A., et al. (2020). Cross-Comparison of Human iPSC Motor Neuron Models of Familial and Sporadic ALS Reveals Early and Convergent Transcriptomic Disease Signatures. Cell Syst.

Iridoy, M.O., Zubiri, I., Zelaya, M.V., Martinez, L., Ausin, K., Lachen-Montes, M., Santamaria, E., Fernandez-Irigoyen, J., and Jerico, I. (2018). Neuroanatomical Quantitative Proteomics Reveals Common Pathogenic Biological Routes between Amyotrophic Lateral Sclerosis (ALS) and Frontotemporal Dementia (FTD). Int J Mol Sci 20.

Kelly, R.T. (2020). Single-cell Proteomics: Progress and Prospects. Mol Cell Proteomics 19, 1739–1748.

Klim, J.R., Williams, L.A., Limone, F., Guerra San Juan, I., Davis-Dusenbery, B.N., Mordes, D.A., Burberry, A., Steinbaugh, M.J., Gamage, K.K., Kirchner, R., et al. (2019). ALS-implicated protein TDP-43 sustains levels of STMN2, a mediator of motor neuron growth and repair. Nat Neurosci 22, 167–179.

Krach, F., Batra, R., Wheeler, E.C., Vu, A.Q., Wang, R., Hutt, K., Rabin, S.J., Baughn, M.W., Libby, R.T., Diaz-Garcia, S., et al. (2018). Transcriptome-pathology correlation identifies interplay between TDP-43 and the expression of its kinase CK1E in sporadic ALS. Acta Neuropathol 136, 405–423.

Ladd, A.C., Brohawn, D.G., Thomas, R.R., Keeney, P.M., Berr, S.S., Khan, S.M., Portell, F.R., Shakenov, M.Z., Antkowiak, P.F., Kundu, B., et al. (2017). RNA-seq analyses reveal that cervical spinal cords and anterior motor neurons from amyotrophic lateral sclerosis subjects show reduced expression of mitochondrial DNA-encoded respiratory genes, and rhTFAM may correct this respiratory deficiency. Brain Res 1667, 74–83.

Lee, T., Li, Y.R., Ingre, C., Weber, M., Grehl, T., Gredal, O., de Carvalho, M., Meyer, T., Tysnes, O.B., Auburger, G., et al. (2011). Ataxin-2 intermediate-length polyglutamine expansions in European ALS patients. Hum Mol Genet 20, 1697–1700.

Lehmkuhl, E.M., and Zarnescu, D.C. (2018). Lost in Translation: Evidence for Protein Synthesis Deficits in ALS/FTD and Related Neurodegenerative Diseases. Adv Neurobiol 20, 283–301.

Ling, J.P., Pletnikova, O., Troncoso, J.C., and Wong, P.C. (2015). TDP-43 repression of nonconserved cryptic exons is compromised in ALS-FTD. Science 349, 650–655.

Liu, E.Y., Russ, J., Cali, C.P., Phan, J.M., Amlie-Wolf, A., and Lee, E.B. (2019). Loss of Nuclear TDP-43 Is Associated with Decondensation of LINE Retrotransposons. Cell Rep 27, 1409–1421 e1406.

Liu, G., Coyne, A.N., Pei, F., Vaughan, S., Chaung, M., Zarnescu, D.C., and Buchan, J.R. (2017). Endocytosis regulates TDP-43 toxicity and turnover. Nat Commun 8, 2092.

Lundberg, E., Fagerberg, L., Klevebring, D., Matic, I., Geiger, T., Cox, J., Algenas, C., Lundeberg, J., Mann, M., and Uhlen, M. (2010). Defining the transcriptome and proteome in three functionally different human cell lines. Mol Syst Biol 6, 450.

Ma, X.R., Prudencio, M., Koike, Y., Vatsavayai, S.C., Kim, G., Harbinski, F., Briner, A., Rodriguez, C.M., Guo, C., Akiyama, T., et al. (2022). TDP-43 represses cryptic exon inclusion in the FTD-ALS gene UNC13A. Nature 603, 124–130.

Mackenzie, I.R., and Rademakers, R. (2008). The role of transactive response DNA-binding protein-43 in amyotrophic lateral sclerosis and frontotemporal dementia. Curr Opin Neurol 21, 693–700.

Maier, T., Guell, M., and Serrano, L. (2009). Correlation of mRNA and protein in complex biological samples. FEBS Lett 583, 3966–3973.

Martin-de-Saavedra, M.D., Dos Santos, M., Culotta, L., Varea, O., Spielman, B.P., Parnell, E., Forrest, M.P., Gao, R., Yoon, S., McCoig, E., et al. (2022). Shed CNTNAP2 ectodomain is detectable in CSF and regulates Ca(2+) homeostasis and network synchrony via PMCA2/ATP2B2. Neuron 110, 627–643 e629.

Melamed, Z., Lopez-Erauskin, J., Baughn, M.W., Zhang, O., Drenner, K., Sun, Y., Freyermuth, F., McMahon, M.A., Beccari, M.S., Artates, J.W., et al. (2019). Premature polyadenylation-mediated loss of stathmin-2 is a hallmark of TDP-43-dependent neurodegeneration. Nat Neurosci 22, 180–190.

Miller, R.G., Mitchell, J.D., and Moore, D.H. (2012). Riluzole for amyotrophic lateral sclerosis (ALS)/motor neuron disease (MND). Cochrane Database Syst Rev, CD001447.

Muzio, L., Sirtori, R., Gornati, D., Eleuteri, S., Fossaghi, A., Brancaccio, D., Manzoni, L., Ottoboni, L., Feo, L., Quattrini, A., et al. (2020). Retromer stabilization results in neuroprotection in a model of Amyotrophic Lateral Sclerosis. Nat Commun 11, 3848.

Neumann, M., Sampathu, D.M., Kwong, L.K., Truax, A.C., Micsenyi, M.C., Chou, T.T., Bruce, J., Schuck, T., Grossman, M., Clark, C.M., et al. (2006). Ubiquitinated TDP-43 in frontotemporal lobar degeneration and amyotrophic lateral sclerosis. Science 314, 130–133.

Oeckl, P., Weydt, P., Thal, D.R., Weishaupt, J.H., Ludolph, A.C., and Otto, M. (2020). Proteomics in cerebrospinal fluid and spinal cord suggests UCHL1, MAP2 and GPNMB as biomarkers and underpins importance of transcriptional pathways in amyotrophic lateral sclerosis. Acta Neuropathol 139, 119–134.

Oh, S., Y, J., L, V., Boxer, A., Sockanathan, S., and Na, C.-H. (2021 (preprint)). Discovery of Biomarkers for Amyotrophic Lateral Sclerosis and Frontotemporal Lobar Degeneration From Human Cerebrospinal Fluid Using Mass Spectrometry-Based Proteomics. Research Square Preprint.

Paganoni, S., Macklin, E.A., Hendrix, S., Berry, J.D., Elliott, M.A., Maiser, S., Karam, C., Caress, J.B., Owegi, M.A., Quick, A., et al. (2020). Trial of Sodium Phenylbutyrate-Taurursodiol for Amyotrophic Lateral Sclerosis. N Engl J Med 383, 919–930.

Payne, S.H. (2015). The utility of protein and mRNA correlation. Trends Biochem Sci 40, 1–3.

Petryszak, R., Keays, M., Tang, Y.A., Fonseca, N.A., Barrera, E., Burdett, T., Fullgrabe, A., Fuentes, A.M., Jupp, S., Koskinen, S., et al. (2016). Expression Atlas update--an integrated database of gene and protein expression in humans, animals and plants. Nucleic Acids Res 44, D746–752.

Poliak, S., Gollan, L., Martinez, R., Custer, A., Einheber, S., Salzer, J.L., Trimmer, J.S., Shrager, P., and Peles, E. (1999). Caspr2, a new member of the neurexin superfamily, is localized at the juxtaparanodes of myelinated axons and associates with K+ channels. Neuron 24, 1037–1047.

Poliak, S., Gollan, L., Salomon, D., Berglund, E.O., Ohara, R., Ranscht, B., and Peles, E. (2001). Localization of Caspr2 in myelinated nerves depends on axon-glia interactions and the generation of barriers along the axon. J Neurosci 21, 7568–7575.

Qureshi, Y.H., Berman, D.E., Marsh, S.E., Klein, R.L., Patel, V.M., Simoes, S., Kannan, S., Petsko, G.A., Stevens, B., and Small, S.A. (2022). The neuronal retromer can regulate both neuronal and microglial phenotypes of Alzheimer’s disease. Cell Rep 38, 110262.

R Core Team (2020). R: A language and environment for statistical computing. R Foundation for Statistical Computing, Vienna, Austria.

Rabin, S.J., Kim, J.M., Baughn, M., Libby, R.T., Kim, Y.J., Fan, Y., Libby, R.T., La Spada, A., Stone, B., and Ravits, J. (2010). Sporadic ALS has compartment-specific aberrant exon splicing and altered cell-matrix adhesion biology. Hum Mol Genet 19, 313–328.

Ritchie, M.E., Phipson, B., Wu, D., Hu, Y., Law, C.W., Shi, W., and Smyth, G.K. (2015). limma powers differential expression analyses for RNA-sequencing and microarray studies. Nucleic Acids Res 43, e47.

Scott, R., Sanchez-Aguilera, A., van Elst, K., Lim, L., Dehorter, N., Bae, S.E., Bartolini, G., Peles, E., Kas, M.J.H., Bruining, H., et al. (2019). Loss of Cntnap2 Causes Axonal Excitability Deficits, Developmental Delay in Cortical Myelination, and Abnormal Stereotyped Motor Behavior. Cereb Cortex 29, 586–597.

Shannon, P., Markiel, A., Ozier, O., Baliga, N.S., Wang, J.T., Ramage, D., Amin, N., Schwikowski, B., and Ideker, T. (2003). Cytoscape: a software environment for integrated models of biomolecular interaction networks. Genome Res 13, 2498–2504.

Shao, W., Todd, T.W., Wu, Y., Jones, C.Y., Tong, J., Jansen-West, K., Daughrity, L.M., Park, J., Koike, Y., Kurti, A., et al. (2022). Two FTD-ALS genes converge on the endosomal pathway to induce TDP-43 pathology and degeneration. Science 378, 94–99.

Sharma, K., Schmitt, S., Bergner, C.G., Tyanova, S., Kannaiyan, N., Manrique-Hoyos, N., Kongi, K., Cantuti, L., Hanisch, U.K., Philips, M.A., et al. (2015). Cell type- and brain region-resolved mouse brain proteome. Nat Neurosci 18, 1819–1831.

Shin, J.E., Geisler, S., and DiAntonio, A. (2014). Dynamic regulation of SCG10 in regenerating axons after injury. Exp Neurol 252, 1–11.

Shin, J.E., Miller, B.R., Babetto, E., Cho, Y., Sasaki, Y., Qayum, S., Russler, E.V., Cavalli, V., Milbrandt, J., and DiAntonio, A. (2012). SCG10 is a JNK target in the axonal degeneration pathway. Proc Natl Acad Sci U S A 109, E3696–3705.

Simoes, S., Guo, J., Buitrago, L., Qureshi, Y.H., Feng, X., Kothiya, M., Cortes, E., Patel, V., Kannan, S., Kim, Y.H., et al. (2021). Alzheimer’s vulnerable brain region relies on a distinct retromer core dedicated to endosomal recycling. Cell Rep 37, 110182.

Sreedharan, J., Blair, I.P., Tripathi, V.B., Hu, X., Vance, C., Rogelj, B., Ackerley, S., Durnall, J.C., Williams, K.L., Buratti, E., et al. (2008). TDP-43 mutations in familial and sporadic amyotrophic lateral sclerosis. Science 319, 1668–1672.

Subramaniam, S., Sixt, K.M., Barrow, R., and Snyder, S.H. (2009). Rhes, a striatal specific protein, mediates mutant-huntingtin cytotoxicity. Science 324, 1327–1330.

Szklarczyk, D., Gable, A.L., Lyon, D., Junge, A., Wyder, S., Huerta-Cepas, J., Simonovic, M., Doncheva, N.T., Morris, J.H., Bork, P., et al. (2019). STRING v11: protein-protein association networks with increased coverage, supporting functional discovery in genome-wide experimental datasets. Nucleic Acids Res 47, D607–D613.

Takahashi, M., Kitaura, H., Kakita, A., Kakihana, T., Katsuragi, Y., Onodera, O., Iwakura, Y., Nawa, H., Komatsu, M., and Fujii, M. (2022). USP10 Inhibits Aberrant Cytoplasmic Aggregation of TDP-43 by Promoting Stress Granule Clearance. Mol Cell Biol 42, e0039321.

Tang, F.L., Zhao, L., Zhao, Y., Sun, D., Zhu, X.J., Mei, L., and Xiong, W.C. (2020). Coupling of terminal differentiation deficit with neurodegenerative pathology in Vps35-deficient pyramidal neurons. Cell Death Differ 27, 2099–2116.

Tank, E.M., Figueroa-Romero, C., Hinder, L.M., Bedi, K., Archbold, H.C., Li, X., Weskamp, K., Safren, N., Paez-Colasante, X., Pacut, C., et al. (2018). Abnormal RNA stability in amyotrophic lateral sclerosis. Nat Commun 9, 2845.

Tyanova, S., Temu, T., Sinitcyn, P., Carlson, A., Hein, M.Y., Geiger, T., Mann, M., and Cox, J. (2016). The Perseus computational platform for comprehensive analysis of (prote)omics data. Nat Methods 13, 731–740.

Umoh, M.E., Dammer, E.B., Dai, J., Duong, D.M., Lah, J.J., Levey, A.I., Gearing, M., Glass, J.D., and Seyfried, N.T. (2018). A proteomic network approach across the ALS-FTD disease spectrum resolves clinical phenotypes and genetic vulnerability in human brain. EMBO Mol Med 10, 48–62.

Vagnozzi, A.N., and Pratico, D. (2019). Endosomal sorting and trafficking, the retromer complex and neurodegeneration. Mol Psychiatry 24, 857–868.

Verde, F., Otto, M., and Silani, V. (2021). Neurofilament Light Chain as Biomarker for Amyotrophic Lateral Sclerosis and Frontotemporal Dementia. Front Neurosci 15, 679199.

Vilarino-Guell, C., Wider, C., Ross, O.A., Dachsel, J.C., Kachergus, J.M., Lincoln, S.J., Soto-Ortolaza, A.I., Cobb, S.A., Wilhoite, G.J., Bacon, J.A., et al. (2011). VPS35 mutations in Parkinson disease. Am J Hum Genet 89, 162–167.

Volkening, K., Leystra-Lantz, C., Yang, W., Jaffee, H., and Strong, M.J. (2009). Tar DNA binding protein of 43 kDa (TDP-43), 14-3-3 proteins and copper/zinc superoxide dismutase (SOD1) interact to modulate NFL mRNA stability. Implications for altered RNA processing in amyotrophic lateral sclerosis (ALS). Brain Res 1305, 168–182.

Watanabe, R., Higashi, S., Nonaka, T., Kawakami, I., Oshima, K., Niizato, K., Akiyama, H., Yoshida, M., Hasegawa, M., and Arai, T. (2020). Intracellular dynamics of Ataxin-2 in the human brains with normal and frontotemporal lobar degeneration with TDP-43 inclusions. Acta Neuropathol Commun 8, 176.

Wen, L., Tang, F.L., Hong, Y., Luo, S.W., Wang, C.L., He, W., Shen, C., Jung, J.U., Xiong, F., Lee, D.H., et al. (2011). VPS35 haploinsufficiency increases Alzheimer’s disease neuropathology. J Cell Biol 195, 765–779.

Wickham, H. (2016). ggplot2: Elegant Graphics for Data Analysis (Springer-Verlag New York).

Wilhelm, M., Schlegl, J., Hahne, H., Gholami, A.M., Lieberenz, M., Savitski, M.M., Ziegler, E., Butzmann, L., Gessulat, S., Marx, H., et al. (2014). Mass-spectrometry-based draft of the human proteome. Nature 509, 582–587.

Witzel, S., Maier, A., Steinbach, R., Grosskreutz, J., Koch, J.C., Sarikidi, A., Petri, S., Gunther, R., Wolf, J., Hermann, A., et al. (2022). Safety and Effectiveness of Long-term Intravenous Administration of Edaravone for Treatment of Patients With Amyotrophic Lateral Sclerosis. JAMA Neurol 79, 121–130.

Xue, Y.C., Ng, C.S., Xiang, P., Liu, H., Zhang, K., Mohamud, Y., and Luo, H. (2020). Dysregulation of RNA-Binding Proteins in Amyotrophic Lateral Sclerosis. Front Mol Neurosci 13, 78.

Zhu, Y., Clair, G., Chrisler, W.B., Shen, Y., Zhao, R., Shukla, A.K., Moore, R.J., Misra, R.S., Pryhuber, G.S., Smith, R.D., et al. (2018a). Proteomic Analysis of Single Mammalian Cells Enabled by Microfluidic Nanodroplet Sample Preparation and Ultrasensitive NanoLC-MS. Angew Chem Int Ed Engl 57, 12370–12374.

Zhu, Y., Dou, M., Piehowski, P.D., Liang, Y., Wang, F., Chu, R.K., Chrisler, W.B., Smith, J.N., Schwarz, K.C., Shen, Y., et al. (2018b). Spatially Resolved Proteome Mapping of Laser Capture Microdissected Tissue with Automated Sample Transfer to Nanodroplets. Mol Cell Proteomics 17, 1864–1874.

Zhu, Y., Piehowski, P.D., Zhao, R., Chen, J., Shen, Y., Moore, R.J., Shukla, A.K., Petyuk, V.A., Campbell-Thompson, M., Mathews, C.E., et al. (2018c). Nanodroplet processing platform for deep and quantitative proteome profiling of 10-100 mammalian cells. Nat Commun 9, 882.

Zimprich, A., Benet-Pages, A., Struhal, W., Graf, E., Eck, S.H., Offman, M.N., Haubenberger, D., Spielberger, S., Schulte, E.C., Lichtner, P., et al. (2011). A mutation in VPS35, encoding a subunit of the retromer complex, causes late-onset Parkinson disease. Am J Hum Genet 89, 168–175.

